# Swe1 delays cell cycle progression during impaired mitochondrial inheritance to promote mitochondrial homeostasis

**DOI:** 10.64898/2026.09.18.752582

**Authors:** Lisa Dengler, Igor Kukhtevich, Yue Zhang, Hadar Meyer, Maya Schuldiner, Robert Schneider, Jennifer C. Ewald

## Abstract

Mitochondrial homeostasis is essential for cellular health. To maintain mitochondrial homeostasis in proliferating cells, mitochondrial inheritance to daughter cells must be tightly coordinated with growth and cell division. In budding yeast, mitochondrial inheritance is ensured by active transport into the growing bud. Although inheritance defects are well known to affect cell physiology, the regulatory mechanisms linking mitochondrial inheritance to cell cycle control remain unclear. Here, we use the inheritance defect of cells depleted of the mitochondrial inheritance adaptor Mmr1 to investigate this coordination. Time-resolved analysis of more than 8,000 budding events revealed that budding duration is directly linked to the time of mitochondrial inheritance. To understand the underlying mechanism, we performed a genetic interaction screen using the auxin-inducible degron library. Strains depleted of cell cycle regulators, particularly those involved in mitotic progression, were highly enriched in clones with synthetic growth defects. Among these, Swe1, the bud morphogenesis checkpoint kinase, emerged as a key candidate affecting budding duration in response to impaired mitochondrial inheritance. Co-depletion of Mmr1 and Swe1 strongly reduces the mitochondrial inheritance-dependent lengthening of budding duration, leading to a large fraction of daughter cells with insufficient mitochondrial content. Thus, Swe1-mediated mitotic delay contributes to slowing cell cycle progression in response to defective mitochondrial inheritance. We suggest calling this regulatory loop MIBA – mitochondrial inheritance-dependent budding adaptation. Together, our findings reveal a mechanism by which cells couple mitochondrial inheritance to cell cycle progression and highlight the importance of this coupling for mitochondrial homeostasis across the population.

## Introduction

Mitochondrial homeostasis is critical for cellular health. Mitochondrial content varies across cell types and conditions but is typically maintained at a relatively constant level within one type of cell population (Posakony, England et al. 1977, Rafelski, Viana et al. 2012, Miettinen and Björklund 2016, Marshall 2020, Seel, Padovani et al. 2023). To achieve this, cells need to coordinate mitochondrial inheritance with cell growth during the cell division cycle, ensuring that daughter cells receive a sufficient amount of mitochondria to sustain proliferation and viability. Indeed, reduced mitochondrial concentrations compromise cell growth (Jajoo, Jung et al. 2016, Chacko, Nakaoka et al. 2025, Dengler, Padovani et al. 2026) and impaired mitochondrial function causes cells to arrest in G1 (Gorospe, Carvalho et al. 2023).

In the asymmetrically dividing budding yeast, mitochondrial inheritance is achieved by the active transport of mitochondria into the growing bud (Klecker and Westermann 2020). The transport of mitochondria into the bud occurs along the actin cytoskeleton and is mediated by the class V myosin Myo2. Mitochondrial transport by Myo2 relies on the adaptor proteins Mmr1 and Ypt11 which interact with the cargo-binding domain of Myo2 and the mitochondrial outer membrane (Itoh, Watabe et al. 2002, Itoh, Toh et al. 2004, Frederick, Okamoto et al. 2008, Eves, Jin et al. 2012, Tang, Li et al. 2019). This system ensures that mitochondria are actively transported to the bud at an early phase of bud growth. Deletion of both Mmr1 and Ypt11 was shown to be synthetically lethal in the commonly used lab strain S288C (Chernyakov, Santiago-Tirado et al. 2013) demonstrating the essentiality of the adaptors for mitochondrial inheritance and cell growth.

Mitochondrial inheritance has been shown to be coupled to mitotic spindle positioning and mitotic exit (García-Rodríguez, Crider et al. 2009, Kraft and Lackner 2017). In addition to these specific events, delayed mitochondrial inheritance has been associated with slowed and prolonged bud growth in Ypt11 deletion cells, suggesting a link between mitochondrial inheritance and budding progression (Rafelski, Viana et al. 2012). However, how this adaptation is achieved and which regulators play an important role adjusting budding progression under conditions in which mitochondrial inheritance is impaired, remains unclear.

Here, we investigate the interplay between mitochondrial inheritance, bud growth and cell cycle progression on a single-cell level. Although mitochondrial inheritance defects occur in wild-type cells, their frequency is low. To provoke a strong mitochondrial inheritance defect we therefore used auxin-inducible degradation (AID) (Yesbolatova, Saito et al. 2020) of the adaptor protein Mmr1. Our time-resolved analysis of more than 8,000 budding events revealed that bud growth and budding duration are directly dependent on the dynamics of mitochondrial inheritance. We next performed a genetic screen with a proteome-wide AID library (Valenti, David et al. 2025) to identify negative genetic interactors of Mmr1, which identified the bud morphogenesis checkpoint kinase Swe1. Follow-up work demonstrated that Swe1 serves as a regulator delaying cell cycle progression when mitochondrial inheritance is impaired. More broadly, our work uncovers a regulatory circuit connecting spatial organization of an organelle with biosynthetic capacity and cell cycle control to ensure population level homeostasis.

## Results

### Mmr1 depletion induces a rapid and strong mitochondrial inheritance defect

To investigate the mechanisms underlying the coordination of mitochondrial inheritance with cell cycle progression we exploited the inheritance defect associated with loss of Mmr1. Using the AID2 system (Yesbolatova, Saito et al. 2020) to acutely deplete Mmr1, we exclude long-term adaptations which occur in deletion cells (Hughes, Roberts et al. 2000, Kafri, Bar-Even et al. 2005, Teng, Dayhoff-Brannigan et al. 2013). After addition of the auxin analog 5-Ph-IAA, the degradation of our target protein Mmr1 is achieved almost completely within 15 min (Fig 1A).

**Figure 1.**
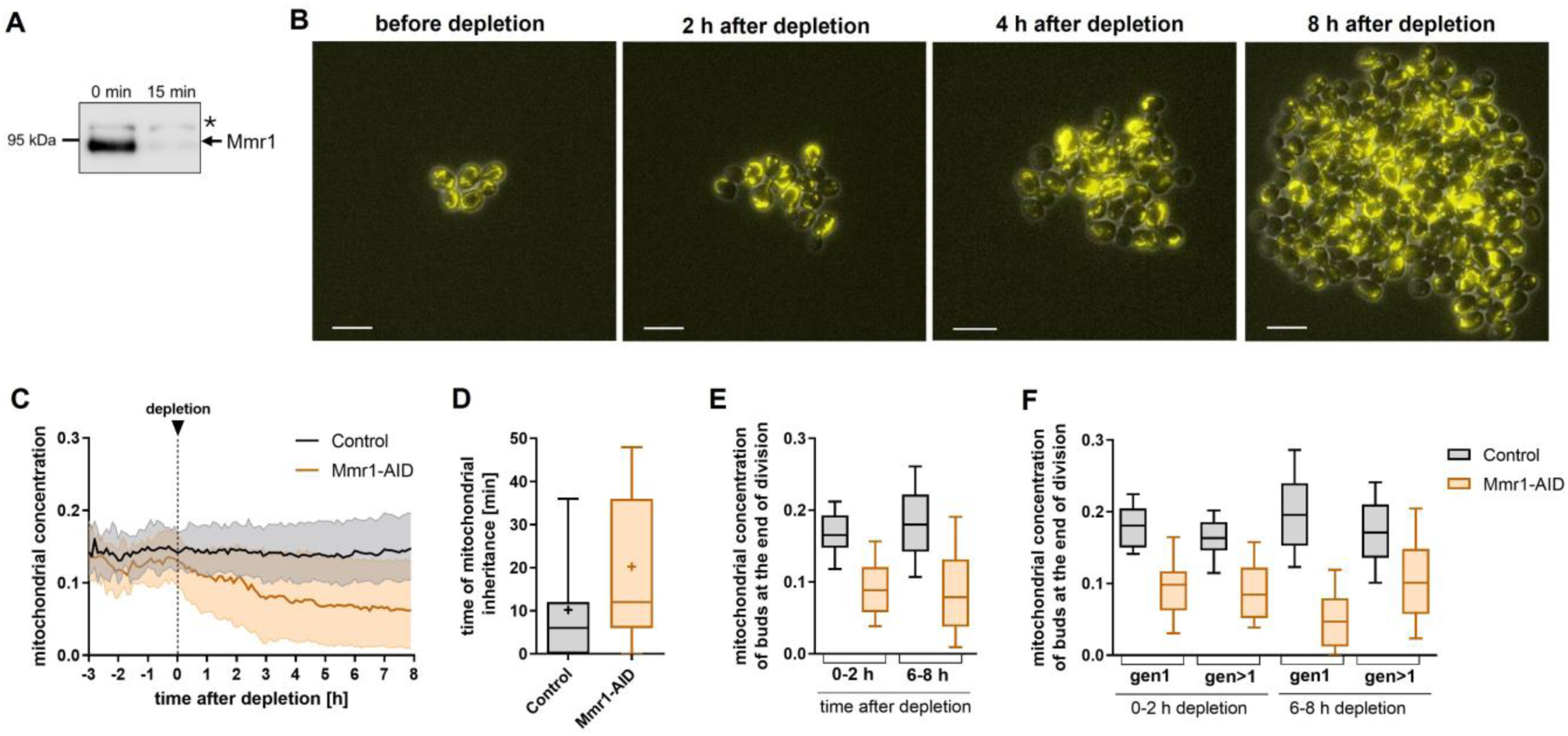
Mmr1 depletion as a tool to study the effects of mitochondrial inheritance. A) – F) Cells were grown in SC-glucose to log phase and 2 µM 5-Ph-IAA was added at 0 h. A) Western blot of the depletion of Mmr1-AID-6xFLAG. * indicates unspecific band. B) Example images of Mmr1-AID cells before and after depletion. Mitochondria are visualized using pre-Su9-mCardinal. Maximum z-projections of mitochondria are shown. Scale bar = 10 µm. C) Mitochondrial concentration (mitochondrial volume/cell volume) over time after Mmr1 depletion from two biological replicates. Medians with 25^th^ and 75^th^ percentiles are shown. 8 h: n (Control) = 3386 cells, n (Mmr1-depleted) = 2533 cells. D) Quantification of the time mitochondria were first detected in the bud after budding started, from two biological replicates. Control: n = 2400, Mmr1-depleted: n = 2174. E) Quantification of the mitochondrial concentration of buds at the end of division from two biological replicates. 0 – 2 h: n (Control) = 86, n (Mmr1-depleted) = 83; 6 – 8 h: n (Control) = 1207, n (Mmr1-depleted) = 822. F) Quantification of the mitochondrial concentration of buds at the end of division in first– and higher-generation divisions from two biological replicates. 0 – 2 h: Control: n (gen 1) = 32, n (gen>1) = 54; Mmr1-depleted: n (gen 1) = 36, n (gen>1) = 47; 6-8 h: Control: n (gen 1) = 518, n (gen>1) = 689; Mmr1-depleted: n (gen 1) = 261, n (gen>1) = 561. D-F Whiskers of box plots indicate 10th and 90th percentiles, line indicates median, where shown + indicates mean, boxes show 25th and 75th percentiles.

To monitor mitochondrial inheritance over the cell cycle, we performed live-cell imaging using the fluorescent protein mCardinal coupled to the mitochondrial targeting sequence pre-Su9, a well-established marker to visualize mitochondria in yeast (Vowinckel, Hartl et al. 2015, Di Bartolomeo, Malina et al. 2020, Dengler, Padovani et al. 2026). Prior to depletion, Mmr1-AID cells growing on SC glucose display wild-type mitochondrial morphology and distribution (Fig 1B), and addition of 5-Ph-IAA alone did not affect mitochondrial morphology and distribution in Control cells (no protein tagged with AID) (Fig. S1A). Depletion of Mmr1 resulted in an immediate and strong mitochondrial inheritance defect already affecting mitochondrial distribution in the first cell cycle after depletion (Fig 1B). Consistent with this observation, the mitochondrial concentration (mitochondrial volume/cell volume) in the population declined rapidly and continuously within the first hours following Mmr1 depletion (Fig 1C).

To quantify the mitochondrial inheritance defect following Mmr1 depletion, we determined the time when mitochondria are first detected in the bud. We monitored budding using Myo1 a as marker, annotated the cell cycle status using Cell-ACDC (Padovani, Mairhörmann et al. 2022) and segmented mitochondria in 3D. We observed that after Mmr1 depletion, buds obtain mitochondria at later stages of budding (Fig 1D). This inheritance delay is in line with previous reports of Δ*mmr1* cells (Itoh, Toh et al. 2004, Förtsch, Hummel et al. 2011, Swayne, Zhou et al. 2011, Chernyakov, Santiago-Tirado et al. 2013), suggesting that acute Mmr1 depletion rapidly induces a phenotype comparable to deletion cells.

We next wanted to determine whether delayed mitochondrial inheritance affects the mitochondrial concentration of buds at the end of division. Before depletion, the mitochondrial concentration of buds at division is comparable between Control and Mmr1-AID cells (Fig. S1B). In contrast, within the first two hours, Mmr1 depletion caused a strong reduction in the mitochondrial concentration of buds and extended depletion decreased the bud mitochondrial concentration further (Fig 1E).

After completing division, buds will become first-generation mothers. Consequently, long-term depletion of Mmr1 results in a reduced mitochondrial concentration of first-generation mothers, whereas higher-generation mothers have higher mitochondrial concentrations than Control mothers (Fig. S1C). We therefore decided to analyze first– and higher-generation cell cycles separately. This generation-resolved analysis revealed that buds of first-generation mothers have a lower mitochondrial concentration at the end of division than buds of higher-generation mothers (Fig 1F).

Taken together, these results demonstrate that acute depletion of Mmr1 induces a rapid and severe mitochondrial inheritance defect, providing a robust system to investigate how defects in mitochondrial inheritance are coordinated with cell cycle progression.

### Delayed mitochondrial transfer is associated with slower and prolonged bud growth

Since Mmr1 depletion increases the frequency of delayed mitochondrial inheritance and reduces mitochondrial accumulation by the end of division, we asked whether the timing of mitochondrial inheritance influences the accumulation of mitochondria in the bud until division. To address this, we grouped cells depending on the time mitochondria were first detected in the bud (“inheritance time”) and followed their mitochondrial volume during budding.

In Control cells, delayed mitochondrial inheritance resulted in a corresponding delay in mitochondrial volume increase (Fig 2A first-generation, and 2D higher-generation). Nevertheless, by the end of division all groups accumulated comparable mitochondrial volumes regardless of their inheritance time (Fig 2C and 2F), indicating that delayed division may act as a compensatory mechanism. Mmr1-depleted cells likewise showed a delay in mitochondrial accumulation with later inheritance (Fig 2B and E). However, buds that inherited mitochondria later accumulated substantially less mitochondrial volume until the end of division than those inheriting them earlier during budding (Fig 2C and 2F, see Fig. S2A for single-cell traces). Mmr1-depleted buds that inherit mitochondria later do not only contain substantially less mitochondrial volume at the end of division than those inheriting them early but also represent a larger fraction of the population compared to Control cells (Fig 2A,B, 2C,D inserts).

**Figure 2.**
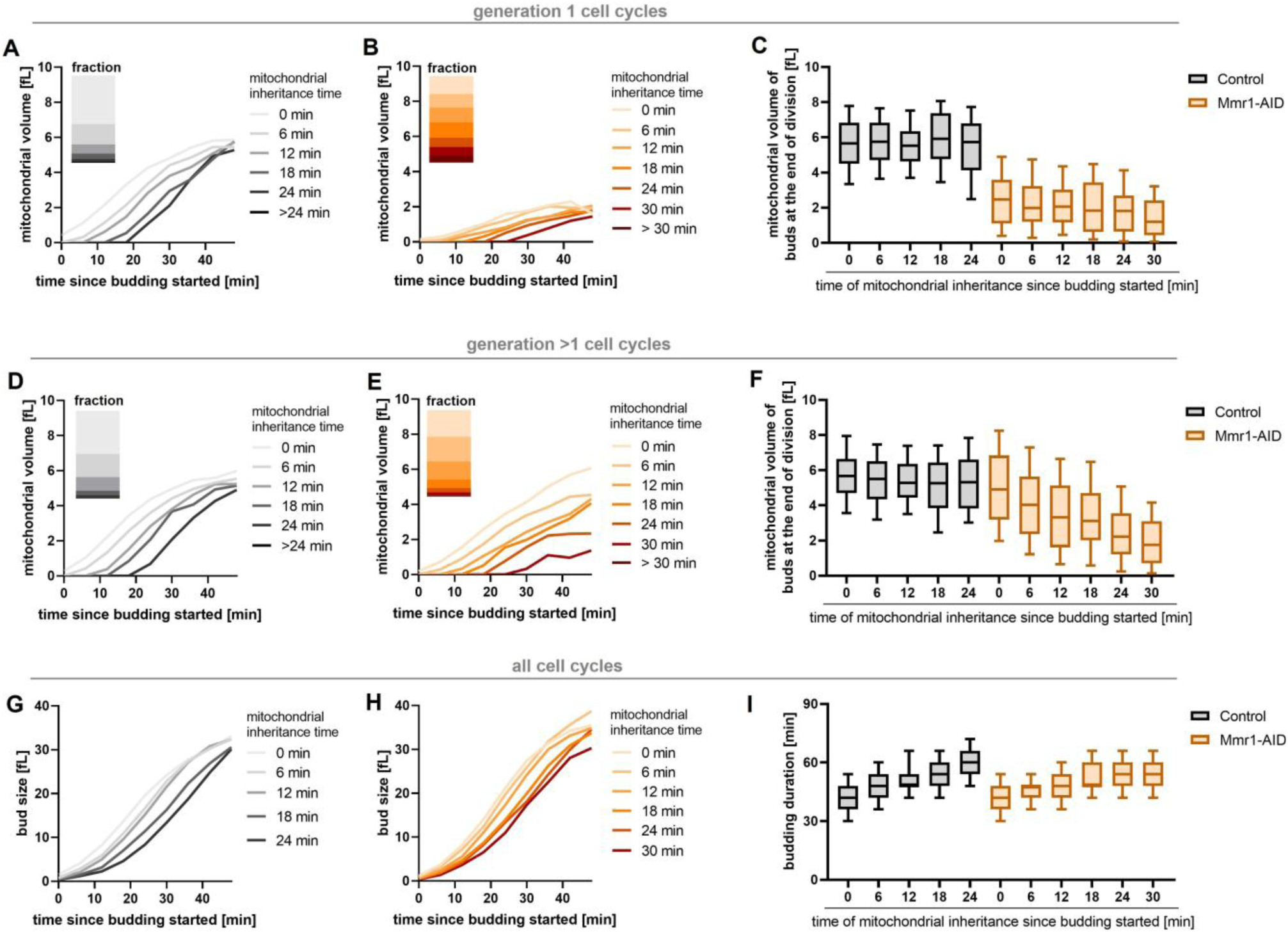
Delayed mitochondrial inheritance results in slower and prolonged bud growth. A) – I) Cells were grown in SC-glucose to log phase and 2 µM 5-Ph-IAA was added to deplete Mmr1. A-C) Analysis of first-generation cell cycles from two biological replicates. Control: n (0 min) = 501, n (6 min) = 210, n (12 min) = 94, n (18 min) = 58, n (24 min) = 30; Mmr1-AID: n (0 min) = 95, n (6 min) = 73, n (12 min) = 78, n (18 min) = 83, n (24 min) = 50, n (30 min) = 47. A) and B) Median mitochondrial volume of the bud depending on the inheritance time of Control A) and Mmr1-AID depleted buds B). Insets show fraction of the population. C) Mitochondrial volume of the bud at the end of division. Whiskers of box plots indicate 10th and 90th percentiles, lines indicate the median, boxes show 25th and 75th percentiles. D-F) Analysis of higher-generation cell cycles from two biological replicates. Control: n (0 min) = 596, n (6 min) = 321, n (12 min) = 190, n (18 min) = 68, n (24 min) = 34; Mmr1-AID: n (0 min) = 339, n (6 min) = 312, n (12 min) = 229, n (18 min) = 109, n (24 min) = 57, n (30 min) = 37. D) and E) Median mitochondrial volume of the bud of different inheritance times of Control D) and Mmr1-AID depleted buds E). Insets show fraction of the population. F) Mitochondrial volume of the bud at the end of division. Whiskers of box plots indicate 10th and 90th percentiles, line indicates median, boxes show 25th and 75th percentiles. G-I) Analysis of all generations from two biological replicates. Control: n (0 min) = 1097, n (6 min) = 531, n (12 min) = 284, n (18 min) = 126, n (24 min) = 64; Mmr1-AID: n (0 min) = 434, n (6 min) = 385, n (12 min) = 307, n (18 min) = 192, n (24 min) = 107, n (30 min) = 84. G) and H) Median bud growth grouped by the time mitochondria are inherited. I) Budding duration grouped by the time mitochondria are inherited. Whiskers of box plots indicate 10th and 90th percentiles, line indicates median, boxes show 25th and 75th percentiles.

Interestingly, late mitochondrial transfer to the bud was associated with a lower mitochondrial concentration in mothers before budding, especially in higher-generation cell cycles (Fig. S2G). Consistent with this observation, first-generation cell cycles, in which mothers start with a lower mitochondrial concentration than higher-generation mothers, also showed a higher frequency of delayed inheritance.

After Mmr1 depletion first-generation cells, which display a strong mitochondrial inheritance delay, also exhibit a substantial elongation of the budding duration (Fig. S2B). Using our time-resolved categorization of mitochondrial inheritance of more than two thousand cell cycles, we resolved a continuous dependency of bud growth on mitochondrial inheritance timing (Fig 2G,H, see S2C-F for generation resolved analysis): later mitochondrial inheritance is associated with progressively slower bud growth and extended budding durations in both Control and Mmr1 depleted cells (Fig 2I, see Fig. S2H for generation resolved analysis). The reduced growth rates of the bud are compensated for by the prolonged budding period resulting in comparable final bud sizes across inheritance categories (Fig. S2I).

Taken together, our time-resolved analysis of mitochondrial inheritance shows that delayed mitochondrial inheritance is associated with slower and prolonged bud growth. While Mmr1-depleted cells are not able to fully compensate for delayed mitochondrial inheritance, lengthening of budding duration in response to delayed inheritance prevents a more severe reduction of the mitochondrial volume in buds. In Control cells, this compensatory response maintains a constant mitochondrial volume at the end of division, regardless of when mitochondria are inherited ensuring proper mitochondrial concentrations in the emerging daughter.

### A genetic interaction screen of mitochondrial inheritance defects against a protein-depletion library identifies cell-cycle candidates

Having established that mitochondrial inheritance is coordinated with bud growth and cell cycle progression, we next sought to identify the molecular mechanisms supporting this coordination. To identify factors involved in the cellular response to impaired mitochondrial inheritance, we integrated the Mmr1-AID trait into the recently established AID2-GFP library (Valenti, David et al. 2025) creating double-depletion strains for the majority of the yeast proteome. We then screened the resulting double-AID strains for synthetic growth defects upon co-depletion. The use of the AID system allowed us to rule out secondary defects and changes through adaptations which often occur in knockouts (Schuldiner, Collins et al. 2005, Lengefeld, Hotz et al. 2017). We assumed that any disturbance of the mechanism coordinating cell cycle progression with mitochondrial inheritance likely results in an increased frequency of cells with an insufficient amount of mitochondria and therefore a reduced colony growth (Fig 3A).

**Figure 3.**
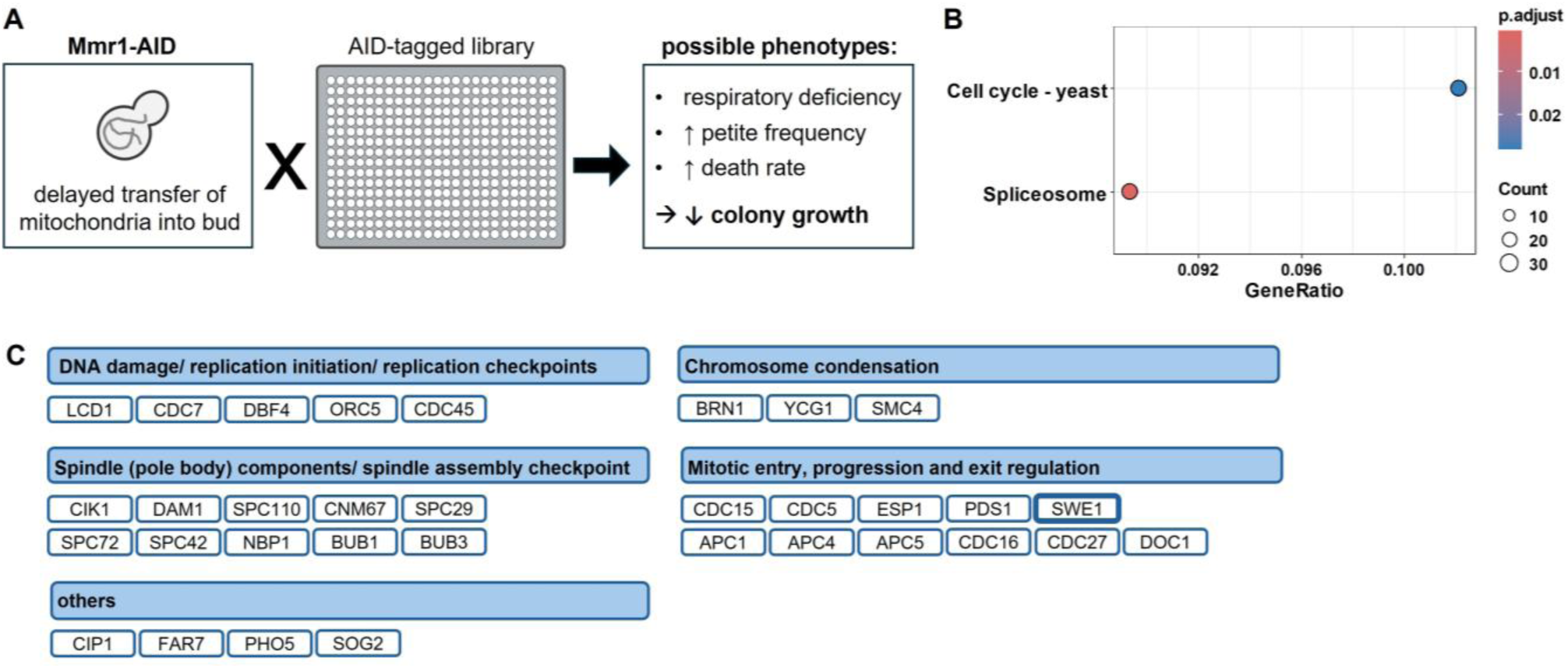
Screen to identify candidates involved in adjusting cell cycle progression in response to mitochondrial inheritance. A) Schematic depiction of the screen and the expected phenotypes. Mmr1-AID-FLAG was integrated into the background of the AID2 library as described in (Valenti et al., 2025) and colony sizes were determined. A negative genetic interaction was scored if the colony size of the double depleted strain was <0.8 compared to both the uninduced strain and the induced single depletion. B) KEGG enrichment analysis of negative genetic interactors with a p-value < 0.05. C) Grouping of cell cycle proteins from the screen.

In support of the validity of this approach, it identified known factors involved in mitochondrial inheritance by strong negative genetic interactions (also previously shown to negatively interact with Mmr1) (Fig. S3C,D): These included Ypt11 and Mdm12 which are both directly involved in mitochondrial inheritance and morphology (Förtsch, Hummel et al. 2011), as well as the phosphatase Ptc1 and its adaptor protein Nbp2 which were both previously associated with mitochondrial inheritance defects (Roeder, Hermann et al. 1998, Jin, Taylor Eves et al. 2009, Hruby, Zapatka et al. 2011, Swayne, Zhou et al. 2011) (Fig. S3C).

To gain more insight into which cellular processes are associated with defective mitochondrial inheritance, we performed KEGG enrichment analysis (Kanehisa and Goto 2000, Kanehisa 2019, Kanehisa, Furumichi et al. 2025). We applied a permissive cutoff of 0.8 of the colony size ratio for both: the double-depleted strains compared to the respective undepleted and single-depleted strains (see legend of Fig. S3A and methods for further explanation). This analysis revealed a significant overrepresentation of spliceosome– and cell cycle-associated proteins (p<0.05) amongst the synthetic genetic interactors (Fig 3B, see Table S1 and S2). Similar results were obtained with a stricter cutoff of 0.7 (Fig. 3B, S3B, Table S1) supporting the robustness of the enrichment. Given our earlier observations linking mitochondrial inheritance defects to altered bud duration, the enrichment of cell cycle regulators was particularly striking and reinforced our focus on cell cycle-associated candidates.

Several of the candidates that showed a defect in our screen were previously shown to affect mitochondrial function and mtDNA abundance (Table S3) (Dimmer, Fritz et al. 2002, Luban, Beutel et al. 2005, Merz and Westermann 2009, Puddu, Herzog et al. 2019, Göke, Schrott et al. 2020, Stenger, Le et al. 2020). To understand at which stages of the cell division cycle co-regulation with mitochondrial inheritance occurs, we grouped the candidate proteins into four functional categories based on their primary biological roles: (1) DNA damage response and replication checkpoint regulators, (2) chromosome condensation, (3) spindle (pole body) and checkpoint proteins, and (4) regulators of mitotic progression and exit (Fig 3C, Table S1).

Given our initial observation linking delayed mitochondrial inheritance to prolonged budding, we got particularly interested in Swe1, a kinase involved in the bud morphogenesis checkpoint. Swe1 participates in regulating mitotic entry by phosphorylating Y19 of Cdk1 to delay cell cycle progression. Cell cycle delay in response to bud formation defects upon actin depolymerization was shown to be mediated by Swe1 (Sia, Herald et al. 1996). This checkpoint activation depends on bud size (Harvey and Kellogg 2003) making Swe1 a compelling candidate for affecting mitotic entry in response to slowed growth during delayed mitochondrial inheritance.

### Loss of Swe1 in the Mmr1 depletion reduces cell size and increases the G1 fraction

To further analyze the genetic interaction between Mmr1 and Swe1, we generated single and double depletion strains *de novo*, confirmed that both proteins are degraded efficiently (Fig 4A), and investigated their interaction at the population level. To understand if Swe1 might affect cell cycle progression in response to impaired mitochondrial inheritance, we measured the cell size of populations growing in SC-glucose. Before depletion, cell size distributions do not differ between the strains (Fig. S4A), but after 6 h a slight shift to smaller cells can be seen (Fig 4B, right).

**Figure 4.**
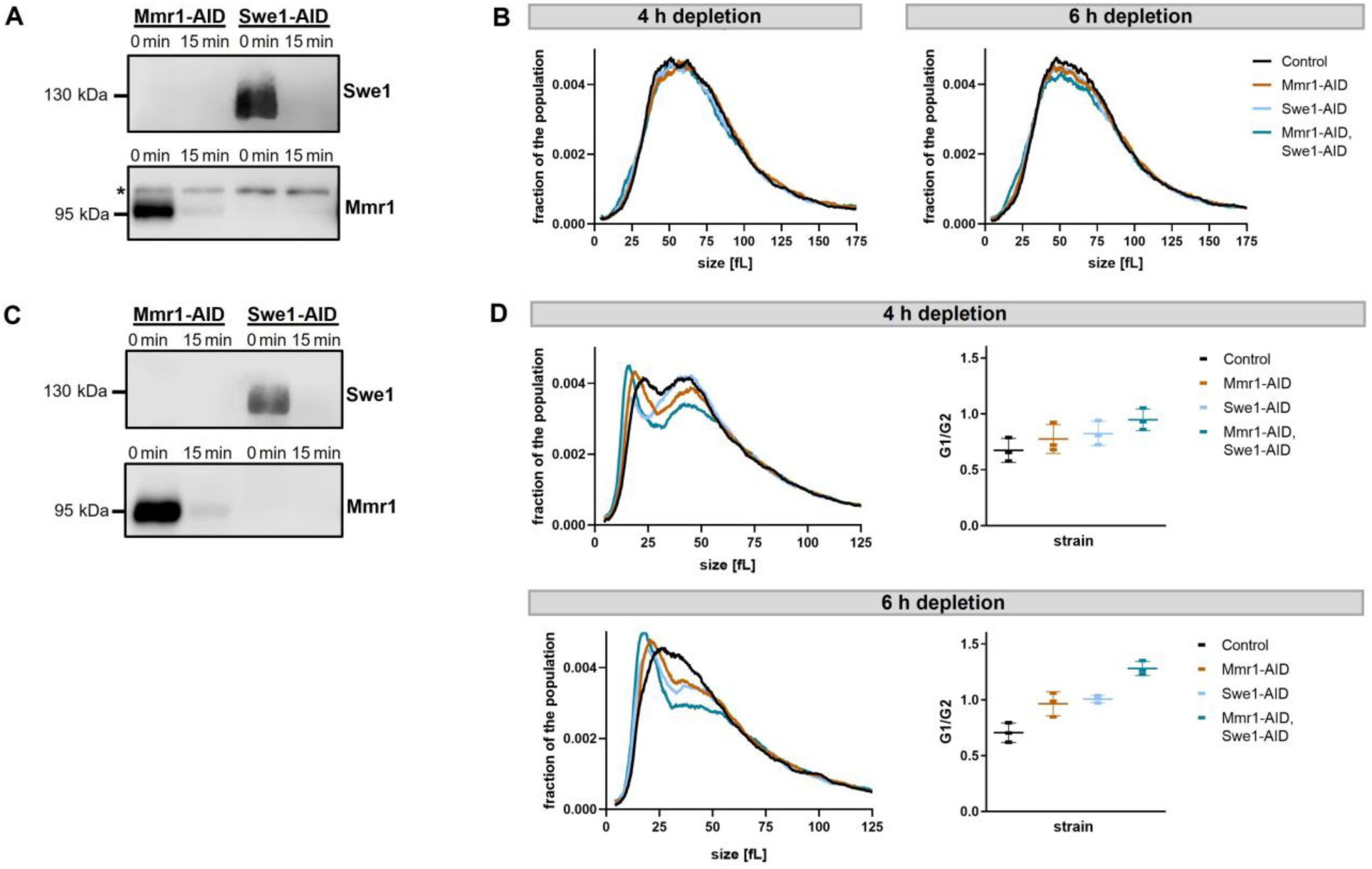
Double-depletion of Mmr1 and Swe1 affects cell size and cell cycle distribution under respiratory conditions. A) Western blot of the depletion of Mmr1-AID-6xFLAG and Swe1-AID-6xFLAG of cells grown in SC-glucose. * indicates unspecific band. B) Cell size distribution of cells grown in SC-glucose 4 h and 6 h after depletion of Mmr1, Swe1, or both using an impedance-based particle counter. The mean from three biological replicates is shown. C) Western blot of the depletion of Mmr1-AID-6xFLAG and Swe1-AID-6xFLAG of cells grown in SC-ethanol. D) Left: Cell size distribution of cells grown in SC-ethanol 4 h and 6 h after depletion of Mmr1, Swe1, or both. The mean from four biological replicates is shown. Right: Ratio of cells in G1 and G2 measured by flow cytometry of SYBR green-stained cells. Mean and SD from three biological replicates are shown.

On fermentable carbon sources such as glucose, cells are able to grow even with a low mitochondrial content. However, under respiratory conditions, cells rely more on mitochondrial activity and thus have to closely regulate cell cycle and growth with mitochondrial homeostasis to survive. We therefore performed the same experiments in SC-ethanol where cells rely on respiration. Also in this growth medium, Swe1 and Mmr1 are depleted rapidly and efficiently (Fig 4C). Before depletion the cell size distributions are similar between the strains (Fig. S4C). However, 4 h and 6 h after depletion, the cell size distribution shifts to smaller sizes for the single Mmr1 and single Swe1 depletion but even more strongly for cells depleted for both proteins (Fig 4D, left). This indicates that especially in the double depletion more and / or smaller daughter cells accumulate. To further examine if the cell cycle distribution is shifted through depletion of both Swe1 and Mmr1, we measured DNA content using SYBR staining followed by flow cytometry. While both single depletions exhibit a slight increase in G1 cells, the double depletion exhibits a much stronger increase in G1 cells (Fig 4D, right, S4E,F).

Taken together, loss of Swe1 in cells having a mitochondrial inheritance defect results in a decreased cell size on population level through accumulation of small daughter cells suggesting that Swe1 plays a role in regulating budding duration in response to impaired mitochondrial inheritance, warranting closer investigation on the single-cell level.

### Loss of Swe1 aggravates the mitochondrial distribution phenotype of Mmr1-depleted cells

To determine whether loss of Swe1 affects budding duration when mitochondrial inheritance is impaired, we performed live-cell imaging to follow bud growth, duration and mitochondrial inheritance on SC-ethanol. (Fig. S5A,B). Following Swe1 depletion, budding duration was significantly shortened and buds completed division at a smaller size, whereas mother cell size remained unchanged (Fig. S5E,F) in line with previous reports on Swe1 deletions (Harvey, Charlet et al. 2005). To assess whether these changes affect mitochondrial distribution, we quantified the mitochondrial concentration across the population over time. In a Control background, loss of Swe1 had no detectable effect on median mitochondrial concentration (Fig 5A, left panel). In contrast, when mitochondrial inheritance was impaired through depletion of Mmr1, loss of Swe1 caused a pronounced further reduction in the mitochondrial concentration (Fig 5A, right panel).

**Figure 5.**
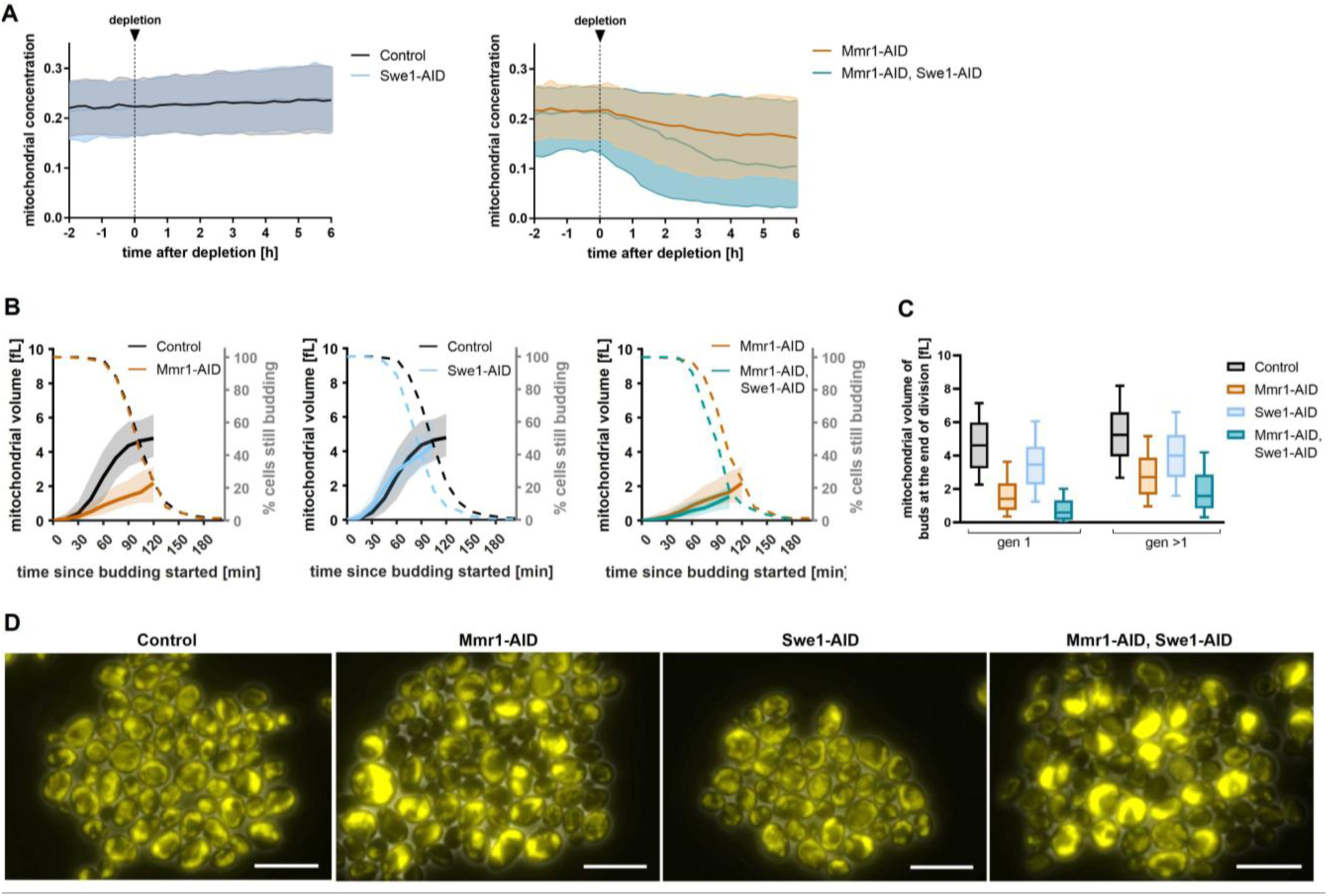
Loss of Swe1 aggravates the mitochondrial distribution phenotype of Mmr1-depleted cells. A) – D) Cells were grown in SC-ethanol and proteins were depleted through addition of 2 µM 5-Ph-IAA. A) Mitochondrial concentrations over time from four biological replicates. Medians with 25^th^ and 75^th^ percentiles are shown. 6 h depletion: n (Control) = 3609 cells, n (Mmr1-AID = 3157 cells, n (Swe1-AID) = 3566 cells, n (Mmr1-AID, Swe1-AID) = 2731 cells. B) Median mitochondrial volume (left axis, solid lines) and percentage of cells that are still budding (right axis, dashed lines) against the time since budding started. Median with 25^th^ and 75^th^ percentiles are shown for the mitochondrial volume until 85 % of the cells have finished budding. Analysis of first-generation cell cycles from four biological replicates is shown. n (Control) = 393, n (Mmr1-AID) = 250, n (Swe1-AID) = 417, n (Mmr1-AID, Swe1-AID) = 191. C) Mitochondrial volume of the buds at the end of division from four biological replicates. Gen 1: n (Control) = 393, n (Mmr1-AID) = 250, n (Swe1-AID) = 417, n (Mmr1-AID, Swe1-AID) = 191; gen >1: n (Control) = 841, n (Mmr1-AID) = 837, n (Swe1-AID) = 1097, n (Mmr1-AID, Swe1-AID) = 893. Whiskers of box plots indicate 10th and 90th percentiles, line indicates median, boxes show 25th and 75th percentiles. D) Example images 6 h after depletion. Maximum z-projections are shown. Scale bar = 10 µm.

To determine whether this phenotype results from altered mitochondrial inheritance, we quantified mitochondrial volume accumulation as well as the fraction of cells which were still in budded phase at a given time point. As expected, Swe1-depleted cells completed budding earlier than Control cells in both the Control and Mmr1-depleted backgrounds (Fig 5B). Swe1 depletion did not impair mitochondrial accumulation rate. In contrast, Mmr1 depletion strongly affected mitochondrial accumulation during budding (Fig 5B, left panel, see Fig. S5G, left panel for higher-generation buds). Notably, simultaneous depletion of Swe1 did not further delay mitochondrial inheritance in either first-(Fig 5B, right) or higher-generation divisions (Fig. S5G, right).

As a consequence of impaired inheritance, Mmr1-depleted buds contained substantially less mitochondrial volume and exhibited a reduced mitochondrial concentration at completion of division (Fig 5C and D, S5C). Although Swe1-depleted buds accumulated less mitochondrial volume by the end of division, their mitochondrial concentration remained comparable to Control cells because of their smaller size. However, when mitochondrial inheritance is impaired, mitochondrial volume and mitochondrial concentration in the bud are strongly reduced at the end of the cell cycle. Importantly, bud mitochondrial concentration was significantly lower than in cells depleted of Mmr1 alone, in both first– and higher-generation divisions (Fig 5C and D). In contrast to buds, insufficient mitochondrial inheritance results in an increase in the mitochondrial concentration of the mothers after Mmr1 depletion and more strikingly through the depletion of Mmr1 and Swe1 (Fig. S5I).

Together, these results show that the shorter budding duration through loss of Swe1 further reduces the mitochondrial concentration of buds in Mmr1-depleted cells. This in turn raises the question whether Swe1 is contributing to cell cycle lengthening when mitochondrial transfer is delayed.

### Loss of Swe1 prevents inheritance-dependent lengthening of budding duration

When mitochondrial transfer to the bud is delayed, the time between initial mitochondrial inheritance and the completion of division is likely an important determinant of how much mitochondria can accumulate in the bud. We therefore grouped cells by the time they first inherited mitochondria and analyzed the time between mitochondrial inheritance and the end of division.

We found that when mitochondria were inherited early during budding, loss of Swe1 did not affect the remaining time between mitochondrial inheritance and the end of division in Mmr1-depleted cells (Fig 6A; left: first generations, right: higher generations, see Fig. S6B for budding durations). However, when mitochondrial inheritance was delayed, loss of Swe1 reduced the time between mitochondrial inheritance and the completion of division. The later mitochondria were inherited, the stronger this effect became (Fig 6A). Consistent with this, cells inheriting mitochondria early showed largely overlapping kinetics across all genotypes, whereas at later inheritance times Swe1-depleted and double-depleted cells completed budding earlier than Control and Mmr1-depleted cells (Fig. S6A).

**Figure 6.**
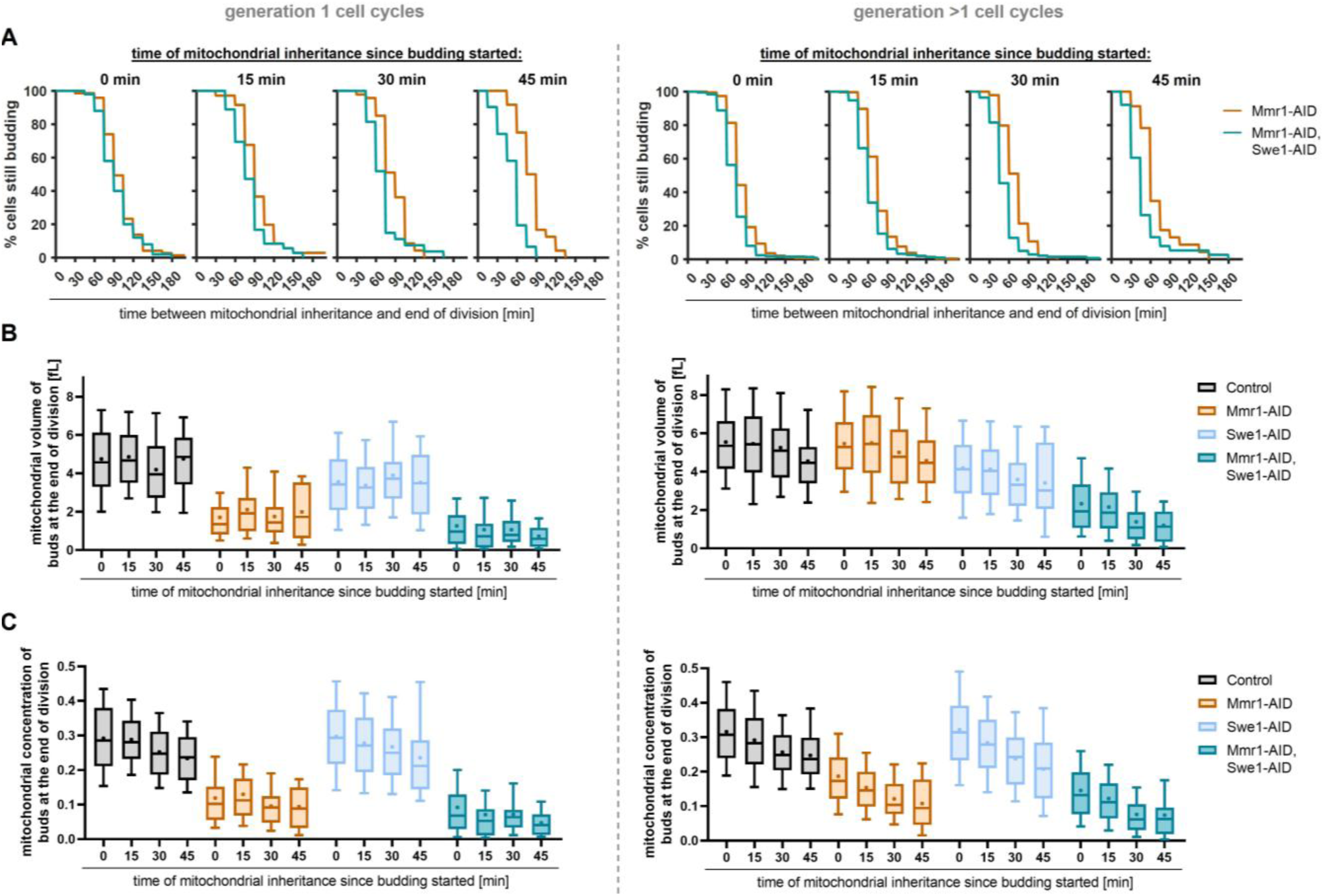
Loss of Swe1 prevents inheritance dependent lengthening of budding duration in Mmr1-depleted cells. A) Time between mitochondrial inheritance and end of division. Left: first-generation, right: higher-generation cell cycles. B) Mitochondrial volume of the buds at the end of division grouped by the time mitochondria are inherited. Analysis of first (left) and higher-generation cells (right) is shown. C) Mitochondrial concentration of the buds at the end of division. Analysis of first (left) and higher-generation cells (right) is shown. Whiskers of box plots indicate 10th and 90th percentiles, line indicates median, + indicates mean, boxes show 25th and 75th percentiles. A) – C) Cells were grown in SC-ethanol and proteins were depleted through addition of 2 µM 5-Ph-IAA. Analysis from four biological replicates is shown. Control (gen 1): n (0 min) = 145, n (15 min) = 96, n (30 min) = 64, n (45 min) = 51; Mmr1-AID (gen 1): n (0 min) = 73, n (15 min) = 71, n (30 min) = 47, n (45 min) = 24; Swe1-AID (gen 1): n (0 min) = 160, n (15 min) = 112, n (30 min) = 84, n (45 min) = 52; Mmr1-AID, Swe1-AID (gen 1): n (0 min) = 50, n (15 min) = 36, n (30 min) = 27, n (45 min) = 31. Control (gen >1): n (0 min) = 390, n (15 min) = 183, n (30 min) = 72, n (45 min) = 27; Mmr1-AID (gen >1): n (0 min) = 491, n (15 min) = 221, n (30 min) = 84, n (45 min) = 35; Swe1-AID (gen >1): n (0 min) = 647, n (15 min) = 316, n (30 min) = 99, n (45 min) = 26; Mmr1-AID, Swe1-AID (gen >1): n (0 min) = 375, n (15 min) = 308, n (30 min) = 141, n (45 min) = 38.

In line with the reduced time between mitochondrial inheritance and division, loss of Swe1 strongly reduced mitochondrial volume and concentration in Mmr1-depleted buds at the end of division (Fig 6B,C and S6D upper panels). The largest effects occurred when mitochondria were inherited late during budding, showing the importance of prolonged budding, especially under conditions of impaired inheritance (Fig. 6B,C and S6D lower panels, see Fig. S6C for bud sizes and S6E for single-cell traces of mitochondrial volumes). Swe1 depletion alone resulted in a slight reduction of the bud mitochondrial concentration at the end of division (Fig 6C and S6D upper panels). This reduction became more prominent with later mitochondrial inheritance (Fig 6C and S6D lower panels). This indicates that extension of budding duration by Swe1 provides additional time for mitochondrial accumulation, which is especially important when mitochondrial inheritance is not only delayed but also impaired.

Taken together, our results show that delayed mitochondrial inheritance is associated with prolonged time cells spend in the budded phases of the cell cycle (S/G2/M). In Control cells, this adaptation allows buds to accumulate similar mitochondrial amounts even in the rare cases when mitochondria are inherited late. Loss of the bud morphogenesis kinase Swe1 disrupts this adaptive response, resulting in a severe impact on mitochondrial homeostasis when mitochondrial inheritance is impaired (Fig 7).

**Figure 7.**
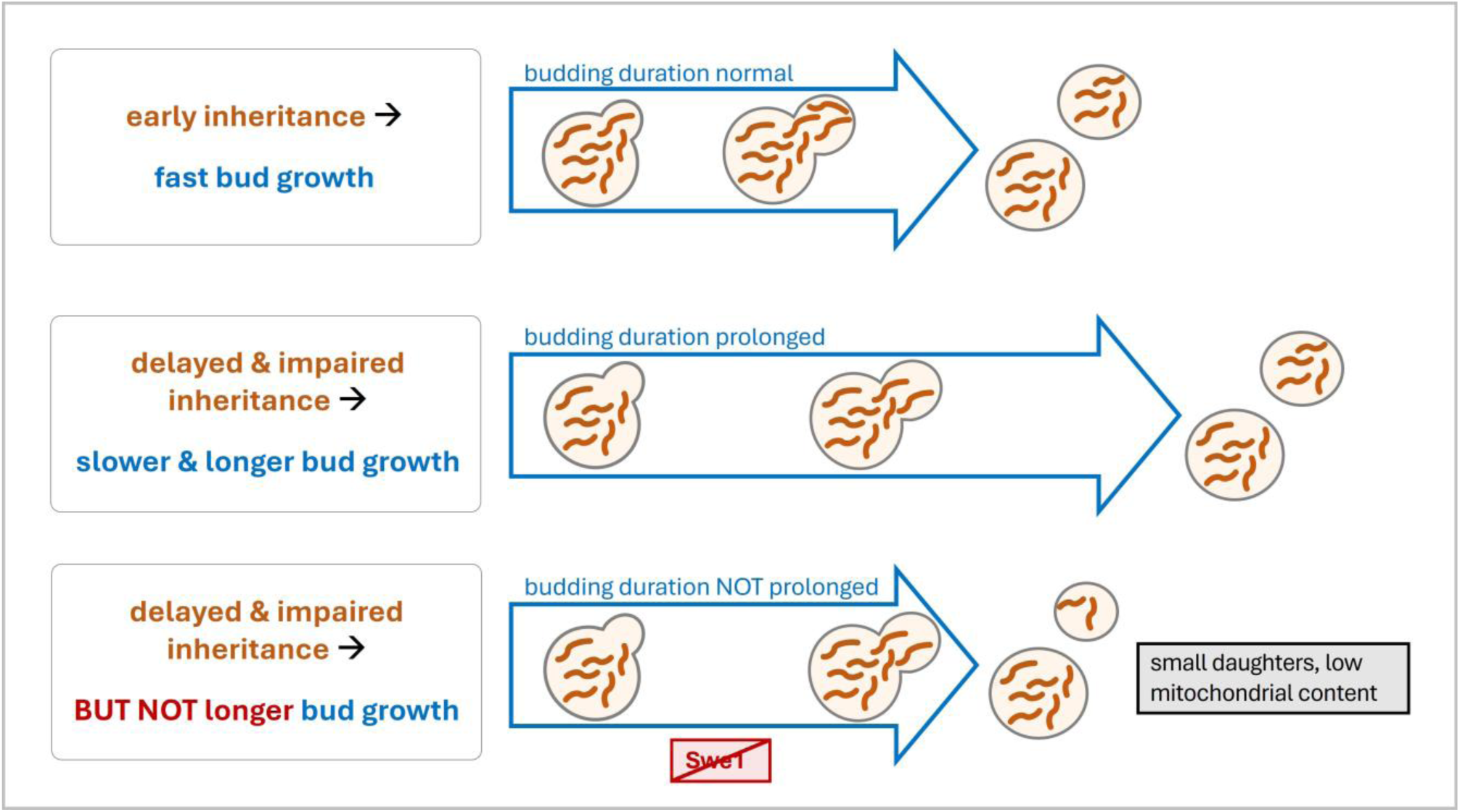
Schematic representation of the suggested role of Swe1 in the regulation of cell cycle progression when mitochondrial inheritance is disturbed. Top: early and sufficient inheritance allows fast bud grow and short budding durations. Middle: delayed mitochondrial inheritance results in slower and longer bud growth. During this, Swe1 likely plays a role in prolonging the budding duration to enable sufficient mitochondrial transfer. Bottom: if mitochondrial transfer is delayed but Swe1 is inactive, budding duration is insufficient and buds obtain an insufficient amount of mitochondria.

## Discussion

### Delayed mitochondrial inheritance progressively slows and prolongs bud growth

In this work, we analyzed more than eight thousand mitochondrial inheritance events to understand how mitochondrial inheritance is coordinated with cell cycle progression. We found that cells adjust the time they spend in the budded phase of the cell cycle according to the timing of mitochondrial inheritance, and that this adaptation becomes particularly important when mitochondrial inheritance is compromised.

Time-resolved analysis revealed a continuous decrease in bud growth rate and a corresponding increase in budding duration with delayed mitochondrial inheritance. In Control cells, this adaptive response maintains stable mitochondrial amounts in the bud independent of the timing of mitochondrial inheritance. This is consistent with previous observations of Ypt11 deletion cells, where delayed bud growth was proposed to allow mitochondrial content to catch up before division (Rafelski, Viana et al. 2012). In Mmr1-depleted cells, however, this adaptive response is insufficient and buds fail to accumulate normal mitochondrial amounts in line with observations made in a recent preprint (Ray, Chen et al. 2025). Nonetheless, prolonged budding prevents a more severe decrease of the mitochondrial concentration upon delayed inheritance in Mmr1 depleted cells. While we induced mitochondrial inheritance defects genetically using the Mmr1-AID depletion, we expect similar effects if cells are exposed to stresses such as increased mitochondrial ROS production which was shown to strongly affect mitochondrial inheritance (Chelius, Bartosch et al. 2023). Likewise, oxidative damage or depolarizing agents may delay efficient transfer of healthy mitochondria to the bud thereby triggering a similar response.

### Genetic screening revealed cell cycle regulators as negative genetic interactors with impaired mitochondrial inheritance

To search for regulators coordinating cell cycle progression with mitochondrial inheritance, we performed the first genome-wide double AID Screen. Screening of negative genetic interactions of Mmr1-AID revealed a significant enrichment of cell cycle genes, supporting the growing evidence of co-regulation between mitochondria and cell cycle regulation (García-Rodríguez, Crider et al. 2009, Crider, García-Rodríguez et al. 2012, Rafelski, Viana et al. 2012, Harbauer, Opalińska et al. 2014, Wang, Fan et al. 2014, Kraft and Lackner 2017, Leite, Costa et al. 2023). Several groups of cell cycle regulators were part of our enrichment, indicating that multiple overlapping mechanisms support successful coordination of mitochondrial inheritance and cell cycle progression. One of the cell cycle groups contained many proteins associated with the mitotic spindle and its assembly checkpoints. Several links between the mitotic spindle and mitochondrial inheritance have been made previously. One link is mediated by Num1 which forms clusters after mitochondrial inheritance that function as cortical attachment sites for dynein. Disrupting mitochondria-driven assembly of Num1 leads to defects in dynein-mediated spindle positioning (Kraft and Lackner 2017). Moreover, several genes which are part of the mitotic spindle or checkpoint proteins of its assembly were previously be found as petite (no mitochondrial DNA, pet) gene/respiratory deficient (Dimmer, Fritz et al. 2002, Luban, Beutel et al. 2005, Merz and Westermann 2009, Puddu, Herzog et al. 2019, Göke, Schrott et al. 2020, Stenger, Le et al. 2020). This is likely to explain the genetic interactions with components of the mitotic spindle and its checkpoint proteins.

In addition to spindle-associated genes, our screen identified many genes involved in mitotic entry, progression and exit regulation as negative genetic interactors of Mmr1 including several components of the anaphase promoting complex (APC/C). Given its central role in regulating mitotic progression through the timely degradation of numerous cell-cycle regulators, this interaction suggests a link between mitochondrial inheritance and mitotic progression. Consistent with this, previous studies have linked the APC/C to mitochondrial inheritance and petite formation (Table S3, (Dutcher 1982, Luban, Beutel et al. 2005, Merz and Westermann 2009, Costanzo, VanderSluis et al. 2016, Puddu, Herzog et al. 2019, Göke, Schrott et al. 2020, Stenger, Le et al. 2020)). Together, our screen identifies several known candidates as well as previously unrecognized connections between mitochondrial inheritance and cell cycle progression particularly in mitosis, and thus suggests further candidates for future mechanistic studies. Based on our observations coupling delayed mitochondrial inheritance with slowed and prolonged budding, we focused on Swe1, a bud morphogenesis checkpoint kinase affecting entry into mitosis.

### Prolonged budding duration in response to delayed mitochondrial inheritance

Swe1 was a particularly interesting candidate among the cell cycle regulators identified in our screen. Cells lacking Swe1 progress more rapidly through G2/M, resulting in a reduced cell volume at birth (Harvey and Kellogg 2003, Soifer and Barkai 2014). Perturbations of bud morphogenesis or actin organization stabilize Swe1 (Sia, Herald et al. 1996). Swe1 in turn phosphorylates Y19-Cdk1 to halt cell cycle progression making Swe1 an interesting possible link in the adaptation of mitochondrial inheritance to cell cycle progression.

At a population level, loss of Swe1 alone causes only minor defects in mitochondrial homeostasis (Fig 6C, S6D). However, buds that inherit mitochondria unusually late are unable to fully adapt if Swe1 is missing. Although these cells represent only a small fraction of the population and therefore do not affect the population much, it is likely to become important under stress. Mitochondrial ROS stress, for example, strongly affects mitochondrial inheritance (Chelius, Bartosch et al. 2023) and may therefore require a more stringent cell-cycle response. Here, we show that Swe1 is an important regulator for the extension of budding duration when inheritance is impaired. Loss of Swe1 in Mmr1-depleted cells results in a drastic reduction of the mitochondrial concentration in buds especially when mitochondrial inheritance is strongly delayed.

The signal linking impaired mitochondrial inheritance to Swe1 activation remains to be further explored. One likely possibility is that Swe1 responds to bud size or bud growth rate as proposed for the classical bud morphogenesis checkpoint (Lew 2003). Growth rate of buds in turn may directly depend on local mitochondrial activity, which provides precursors, energy, ions, and signaling molecules to the growing bud. Thus, slowed bud growth might be a direct metabolic consequence of insufficient supply from mitochondria, thereby connecting mitochondrial inheritance to the bud-morphogenesis checkpoint and mitotic entry. We propose to call this regulatory loop MIBA – mitochondrial inheritance-dependent budding adaptation.

More broadly – our work identifies a powerful homeostatic mechanism that connects spatial organization of an organelle to biosynthetic capacity and cell cycle progression. Our screen suggests that more regulatory links exist, offering new avenues to explore organelle control over growth and division.

## Material and Methods

### Strain construction

All strains were haploid BY4741 or BY4742 derivatives, see genotype in Appendix Table S4. Strains were constructed using standard PCR-based homologous recombination or integration of linearized plasmids. See Appendix Table S5 for details of plasmids used. All strains constructed in this study are available from the corresponding author upon request.

### Cultivation conditions

Cultures were grown to log phase in SC-glucose (1.7 g/l yeast nitrogen base without amino acids (US Biological), 5 g/l ammonium sulfate, 50 mM potassium phthalate, pH adjusted to 5 with KOH, synthetic complete with all amino acids, see Appendix Table S6 for concentrations). Cultures were grown to log phase (OD_600_ 0.6-1.0) and diluted to OD_600_ = 0.1 (glucose) or 0.2 (ethanol) and depletion was induced by addition of 5-Ph-IAA (2-(5-Phenyl-1H-indol-3-yl)acetic acid, 100 mM Stock, dissolved in DMSO) to a final concentration of 2 µM. OD_600_ was determined every hour over 6 h for SC-glucose and over 10 h for SC-ethanol. Growth rates were determined excluding the first hour after depletion.

### Creation of tailored made AID2 libraries for screening

Introduction of Mmr1-AID-6XFLAG into the AID2 library (Valenti, David et al. 2025) was performed as described previously (Cohen and Schuldiner 2011).

### Hit selection and KEGG enrichment analysis

Hit selection was based on the lowest 5 % and 10 % (for 0.7 and 0.8 cutoff, respectively) from the distribution of normalized colony size of all undepleted strains. The cutoff of 0.7 or 0.8 (see Table S9 for full list of hits) was applied for both comparisons of each gene: double-depleted vs undepleted and double-depleted vs single-depleted. KEGG pathway enrichment analysis (Kanehisa and Goto 2000, Kanehisa 2019, Kanehisa, Furumichi et al. 2025) was performed in R (Version 4.4.3) using the Bioconductor package *clusterProfiler*. Gene identifiers were converted from Ensembl to Entrez format using the org.Sc.sgd.db annotation package. Enriched pathways for *Saccharomyces cerevisiae* (organism code: “sce”) were identified with the enrichKEGG() function using a p-value of 0.05.

### Microfluidic cultivation and microscopy for SC-glucose experiments

In preparation for live cell imaging, cells were grown over night in SC-glucose, diluted 1:50 the next morning and grown for 6 h. Cells were sonicated at low power for 3 s and loaded onto a commercial microfluidics system (Y04C-02 plates, CellASIC ONIX2 system, Merck). SC-glucose medium was supplied with a pressure of 3 psi. Cells were grown inside the microfluidic chamber for the duration of 1 h before imaging started. The temperature was kept constant at 30°C using an incubator chamber surrounding the imaging system (Okolab Cage Incubator, Okolab USA INC, San Bruno, CA). Microscopy was performed on a Nikon Ti2 inverted epifluorescence microscope (Nikon Instruments, Japan) with a Lumencor SPECTRA X light engine (Lumencor, Beaverton, USA), a Photometrics Prime 95 (Teledyne Photometrics, USA) backilluminated sCMOS camera. The system was programmed and controlled by the Nikon software NIS Elements. Focus was maintained using the Nikon “Perfect Focus System.” The time-lapse images were taken using a Nikon PlanApo oil-immersion 100× objective (NA = 1.45) with a frequency of 6 min. 5 z-slices were acquired with a step size of 0.5 µm. See Appendix Table S7 for optical filters and Appendix Table S8 for exposure settings. For all fluorophores and tagged proteins, we checked for absence of phototoxicity (Cuny, Schlottmann et al. 2022) by observing mitochondrial morphology and concentration in Control cells over time (see Fig 1C, 5A and S1A).

### Microfluidic cultivation and microscopy for SC-ethanol experiments

In preparation for live cell imaging, cells were grown for ∼20 h, diluted to OD_600_ 0.06 and grown for another 20 h to reach log phase. For live single-cell experiments, we used a previously published custom-made microfluidic device that allows isolating cells in a dedicated region of interest and limits colony growth to the XY-plane (Kukhtevich, Rivero-Romano et al. 2022). The device has eight separate cell culture chambers with a controllable medium exchange that enables parallel imaging of up to eight strains.

The microfluidic device was fabricated employing standard soft lithography. Briefly, by using photolithography, a master mold for replication of the device design in polydimethylsiloxane (PDMS) was fabricated from SU-8 photoresist (MicroChem, USA) spin-coated on a 3`″` Si wafer. The master mold was then filled with a 10:1 mixture of the base to curing agent of PDMS kit Sylgard 184 (Dow Corning, USA) and left at 60 °C for 4 h to crosslink the PDMS. After that, the PDMS replica was cut and peeled off from the master mold, and inlets and outlets for tubing connections were made using a 1 mm puncher. Finally, the replica was sealed with a coverslip after both were treated in O_2_ plasma.

A Nikon Eclipse Ti-E with SPECTRA X light engine illumination and an Andor iXon Ultra 888 camera were used for epifluorescence microscopy. A plan-apo λ 100x/1.45NA Ph3 oil immersion objective was used to take phase contrast and fluorescence images. Temperature control was achieved by setting both a custom-made heatable insertion and an objective heater to 30 °C. Images were taken with a frequency of 15 min. 5 z-slices were acquired with a step size of 0.5 µm. See Appendix Table S7 for optical filters and Appendix Table S8 for exposure settings

### Image analysis and data processing

Images were recorded at 12-bit gray scale and then converted to tiffs in Cell-ACDC (Padovani, Mairhörmann et al. 2022). Cell segmentation was performed in Cell-ACDC using YeaZ (Dietler, Minder et al. 2020) (minimum area: 10 pixels, minimum solidity 0.5, maximal elongation 3.0). Each time frame was then visually inspected and any segmentation and tracking errors corrected in Cell-ACDC. Cell volumes were determined automatically in Cell-ACDC as described in (Padovani, Mairhörmann et al. 2022). The median background fluorescence (cell-free area) for each image was determined and subtracted from the signal at each time point. Cell cycle states and the mother-bud connections were annotated manually in Cell-ACDC which allows us to extract mother-daughter relationships and generation numbers. Mitochondria were segmented in 3D using SpotMAX (Padovani, Čavka et al. 2024) based on mitochondrially localized preSu9-mCardinal. For pre-processing in SpotMAX we used a Gaussian filter with sigma 0.7 followed by a Sato filter with two sigmas 1.0 and 1.5 to enhance network-like structures (Sato, Nakajima et al. 1998). The pre-processed images were then segmented by SpotMAX using Li thresholding (Li and Lee 1993) in 3D.

### SDS-PAGE and Western Blot

Pellets were frozen in liquid nitrogen and stored at – 70 °C. Lysates were prepared with a FastPrep® shaking three times 40 s at 6 m/s with a 1-minute break in between each cycle using lysis buffer (50 mM Tris-HCl pH 8.0, 150 mM NaCl, 5 mM EDTA, 1 % Tergitol) supplemented with 2 × EDTA-free protease inhibitor (GoldBio) and 2 x phosphatase inhibitor (GoldBio). 30 – 50 µg total protein were loaded on 8 or 10 % SDS polyacrylamide (29:1 Bio-Rad) gels. SDS gels were blotted using a commercial wet transfer system (Bio-Rad). For detection of 3xFLAG anti-FLAG M2 antibody (Sigma, product number: F1804) (1:5000 in milk powder) and anti-mouse IgG (H+L), HRP conjugate (Promega, W4021) were used. Luminescence was imaged on a Licor Odyssey FC.

### Flow cytometry

Cells were grown as described in cultivation conditions. Cells were fixed in 70 % ethanol and incubated over night at 4 °C while shaking. Cells were washed twice with 50 mM Tris-HCl pH 8.0 and then treated with RNase A (1 mg/mL) for 40 min at 37 °C. After two wash steps, cells were treated with Proteinase K (20 mg/mL) for 1 h at 37 °C and the wash steps were repeated. Staining with SYBR Green I (1:1000) was performed for 1 h at room temperature. After sonication for 5 s at 55 % cells were diluted with 50 mM Tris-HCl to achieve about 1000 events per second. Flow Cytometry was performed at the ZMBP Central Facilities (University of Tuebingen) with a CytoFlex (Beckman Coulter), 488 nm excitation, 525/40 emission for Syber Green I. For each measurement about 30 000 cells were measured. Gating was performed as shown in Figure S4F.

### Bulk cell size measurements

Bulk cell size measurements were performed using a Casy TT (SN: TT2QA2564, OMNI Life Science GmbH & Co. KG, Bremen, Germany) using a 60 µm capillary. For separation of individual cells, cells were sonicated at 30 % power for 3 s. Approximately 15 000 cells were measured per timepoint. Normalization was performed from 2-8 µm diameter for cells grown in SC-ethanol, and from 2-10 µm in SC-glucose.

## Author Contributions

LD, JCE and MS conceived and designed the study. LD performed most experiments and data analysis. IK performed microscopy for ethanol experiments. YZ supported microscopy analysis. HM conducted the AID screen. JCE, RS, and MS supervised the projects. LD wrote the original draft of the manuscript. JCE, MS, IK and RS edited the manuscript. All authors reviewed and approved the manuscript.

The Authors declare no conflicts of interest.

## Data Availability

Strains and plasmids are available from the corresponding author. Imaging data will be deposited in the BioImageArchive upon final publication and in the meantime can be obtained from the corresponding author upon reasonable request.

## Acknowledgements

We thank Serge Pelet, Jan Skotheim, and Jörg Stelling labs for plasmids. We gratefully acknowledge Kenneth Berendzen for help with flow cytometry. We gratefully acknowledge Katja Kleemann and Tina Schneider for technical support. We thank Benedikt Westermann for helpful discussions. We thank Kurt Schmoller for helpful discussions and comments on the manuscript. We thank Denise Dewald for critical discussions and proofreading. JCE gratefully acknowledges the Deutsche Forschungsgemeinschaft (DFG)—GRK 2364 MOMbrane (Projektnummer 327043846). RS gratefully acknowledges funding from the Helmholtz Gesellschaft. Research in the MS lab is generously supported by the Knell Family Center. MS is Incumbent of the Dr. Gilbert Omenn and Martha Darling Professorial Chair in Molecular Genetics.

---

**Figure S1.**
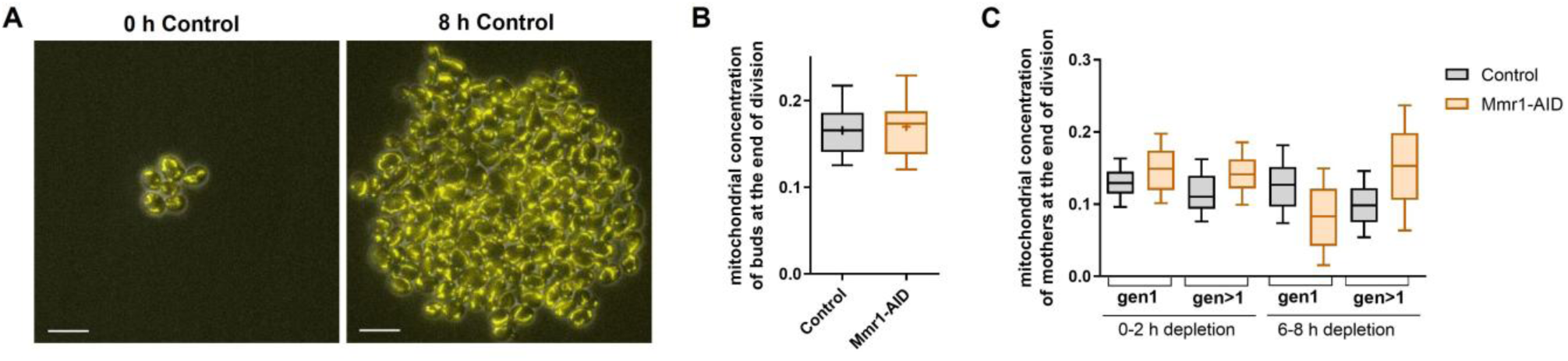
A) – C) Cells were grown in SC-glucose to log phase and 2 µM 5-Ph-IAA was added at 0 h. A) Example images of Control cells before and after addition of 5-Ph-IAA. Mitochondria are visualized using pre-Su9-mCardinal. Maximum z-projections of mitochondria are shown. Scale bar = 10 µm. B) Mitochondrial concentration of buds at the end of division from two biological replicates. Whiskers of box plots indicate 10th and 90th percentiles, line indicates median, + indicates mean, boxes show 25th and 75th percentiles. n (Control) = 31, n (Mmr1-AID) = 34. C) Quantification of the mitochondrial concentration of mothers at the end of division in first– and higher-generation divisions from two biological replicates. 0 – 2 h: Control: n (gen 1) = 32, n (gen>1) = 54; Mmr1-depleted: n (gen 1) = 36, n (gen>1) = 47; 6-8 h: Control: n (gen 1) = 518, n (gen>1) = 689; Mmr1-depleted: n (gen 1) = 261, n (gen>1) = 561. Whiskers of box plots indicate 10th and 90th percentiles, line indicates median, boxes show 25th and 75th percentiles.

**Figure S2.**
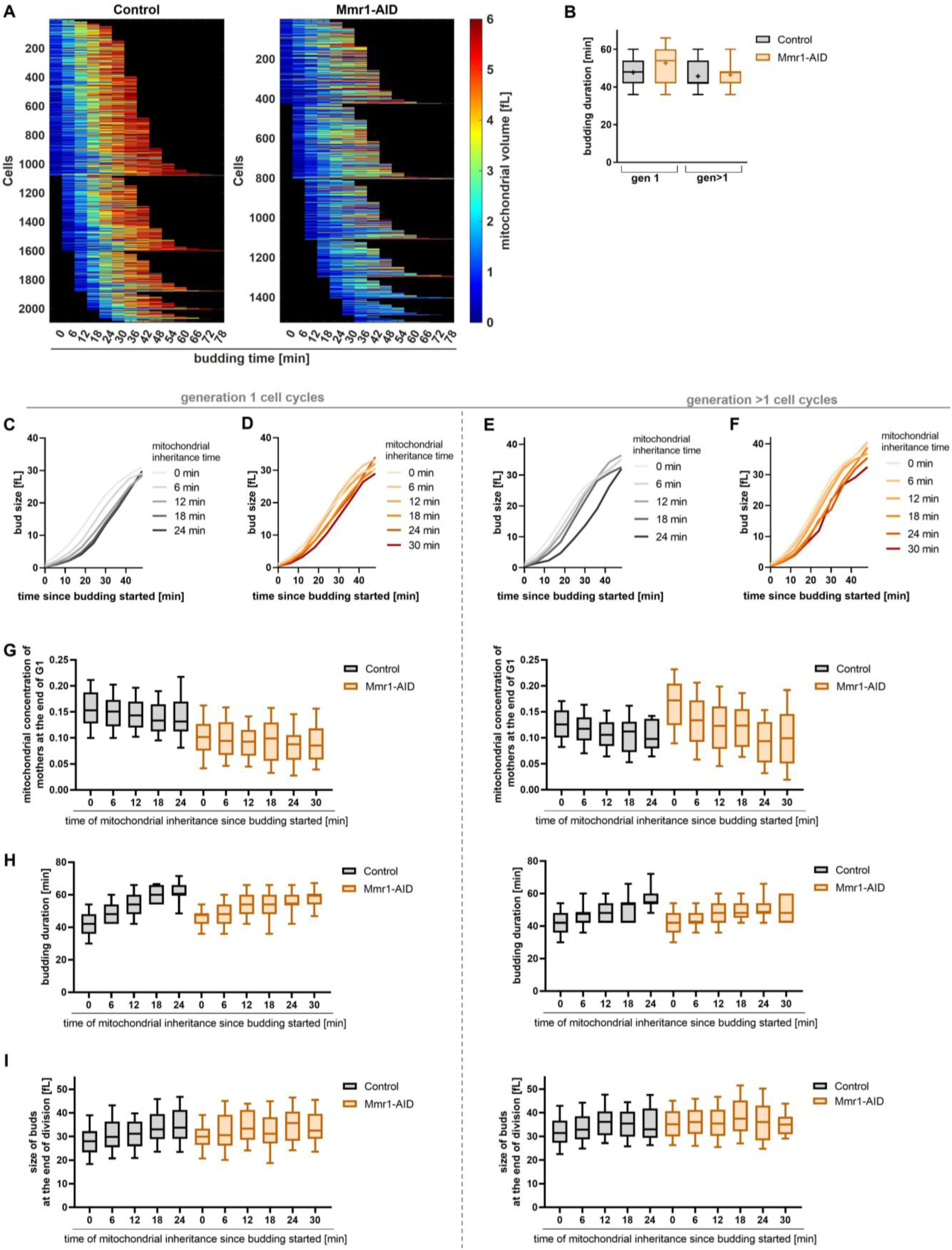
A) – I) Cells were grown in SC-glucose to log phase and 2 µM 5-Ph-IAA was added to deplete Mmr1. A) Heatmaps of the mitochondrial volume of buds through their budding phase from two biological replicates. B) Budding duration for first– and higher-generation divisions from two biological replicates. Whiskers of box plots indicate 10th and 90th percentiles, line indicates median, + indicates mean, boxes show 25th and 75th percentiles. Gen 1: n (Control) = 907, n (Mmr1-AID) = 531; gen >1: n (Control) = 1223, n (Mmr1-AID) = 1114. C-F) Median bud growth grouped by the time mitochondria are inherited. C) and D) Analysis of first-generation cell cycles. C) Control and D) Mmr1-AID depleted buds. E) and F) Analysis of higher-generation cell cycles. E) Control and F) Mmr1-AID depleted buds. G-I) Whiskers of box plots indicate 10th and 90th percentiles, line indicates median, boxes show 25th and 75th percentiles. G) Mitochondrial concentration of mothers at the end of G1, grouped by the time mitochondria are inherited in first (left) and higher-generation (right) cell cycles. H) Budding duration grouped by the time mitochondria are inherited for first (left) and higher-generation (right) cell cycles. I) Bud sizes at the end of division grouped by the time mitochondria are inherited for first (left) and higher-generation (right) cell cycles. C-I) Analysis from two biological replicates is shown. Gen 1 Control: n (0 min) = 501, n (6 min) = 210, n (12 min) = 94, n (18 min) = 58, n (24 min) = 30; gen 1 Mmr1-AID: n (0 min) = 95, n (6 min) = 73, n (12 min) = 78, n (18 min) = 83, n (24 min) = 50, n (30 min) = 47. Gen >1 Control: n (0 min) = 596, n (6 min) = 321, n (12 min) = 190, n (18 min) = 68, n (24 min) = 34; gen >1 Mmr1-AID: n (0 min) = 339, n (6 min) = 312, n (12 min) = 229, n (18 min) = 109, n (24 min) = 57, n (30 min) = 37.

**Figure S3.**
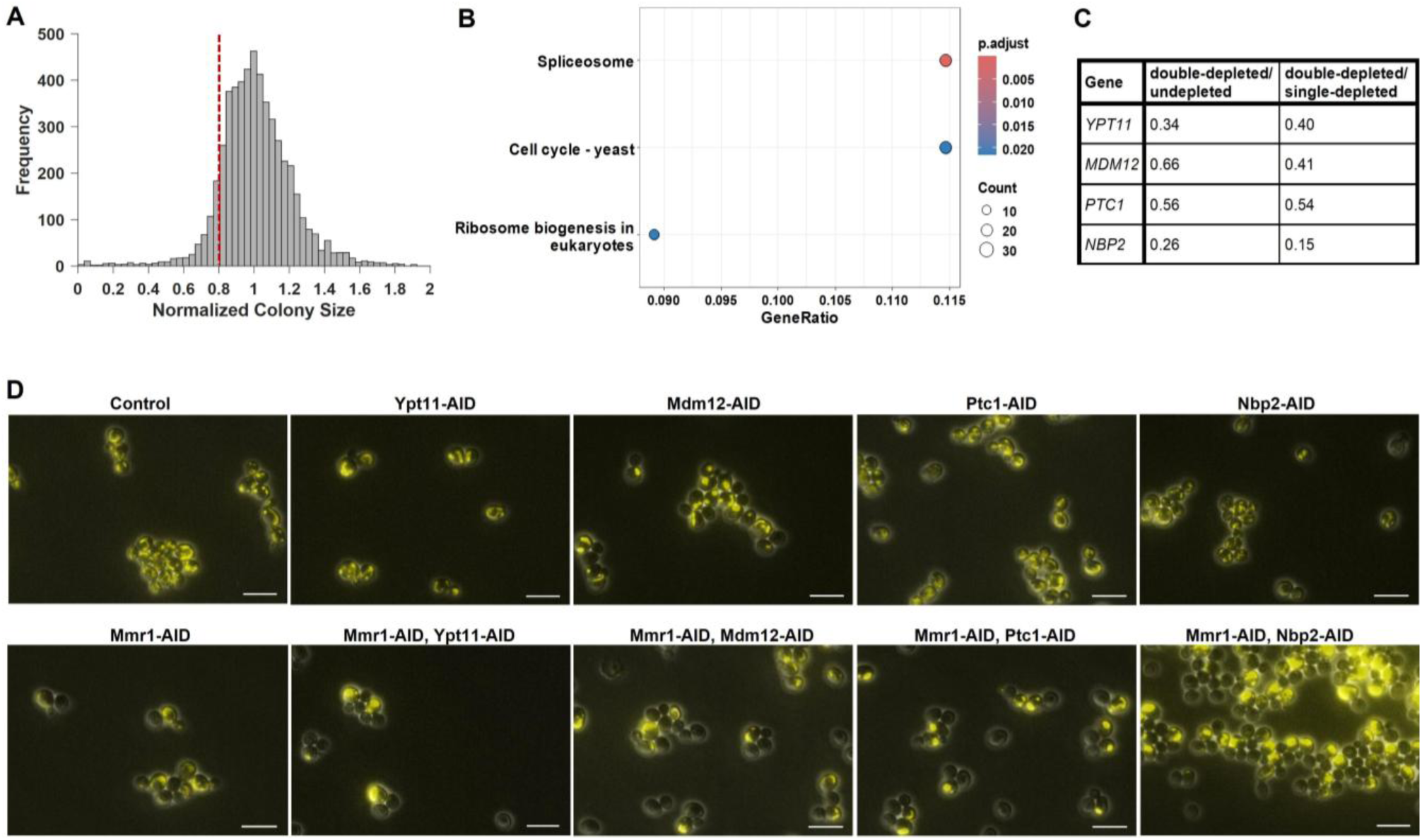
A) Median normalized distribution of the uninduced colony sizes of the double AID-tagged strains, the dashed line depicts the threshold of 0.8. This threshold corresponds to the lowest 10 % of the distribution of all undepleted strains. The stricter threshold of 0.7 corresponds to the lowest 5 % of the distribution of all undepleted strains. These thresholds were used for both comparisons: double-depleted vs undepleted and single-depleted vs double-depleted to select candidate genes. B) KEGG enrichment analysis of negative genetic interactors with a p-value < 0.05. Based on <0.7 relative colony size before depletion and <0.7 relative colony size compared to single AID. C) Ratio of the colony size of doubled-depleted strains vs undepleted and double-depleted vs single-depleted strains. D) Example images of known interactors of mitochondrial inheritance defects identified in the screen. Proteins were depleted for 3 h using 2 µM 5-Ph-IAA. Maximum z-projections of pre-Su9-mCardinal are shown. Scale bar = 10 µm.

**Figure S4.**
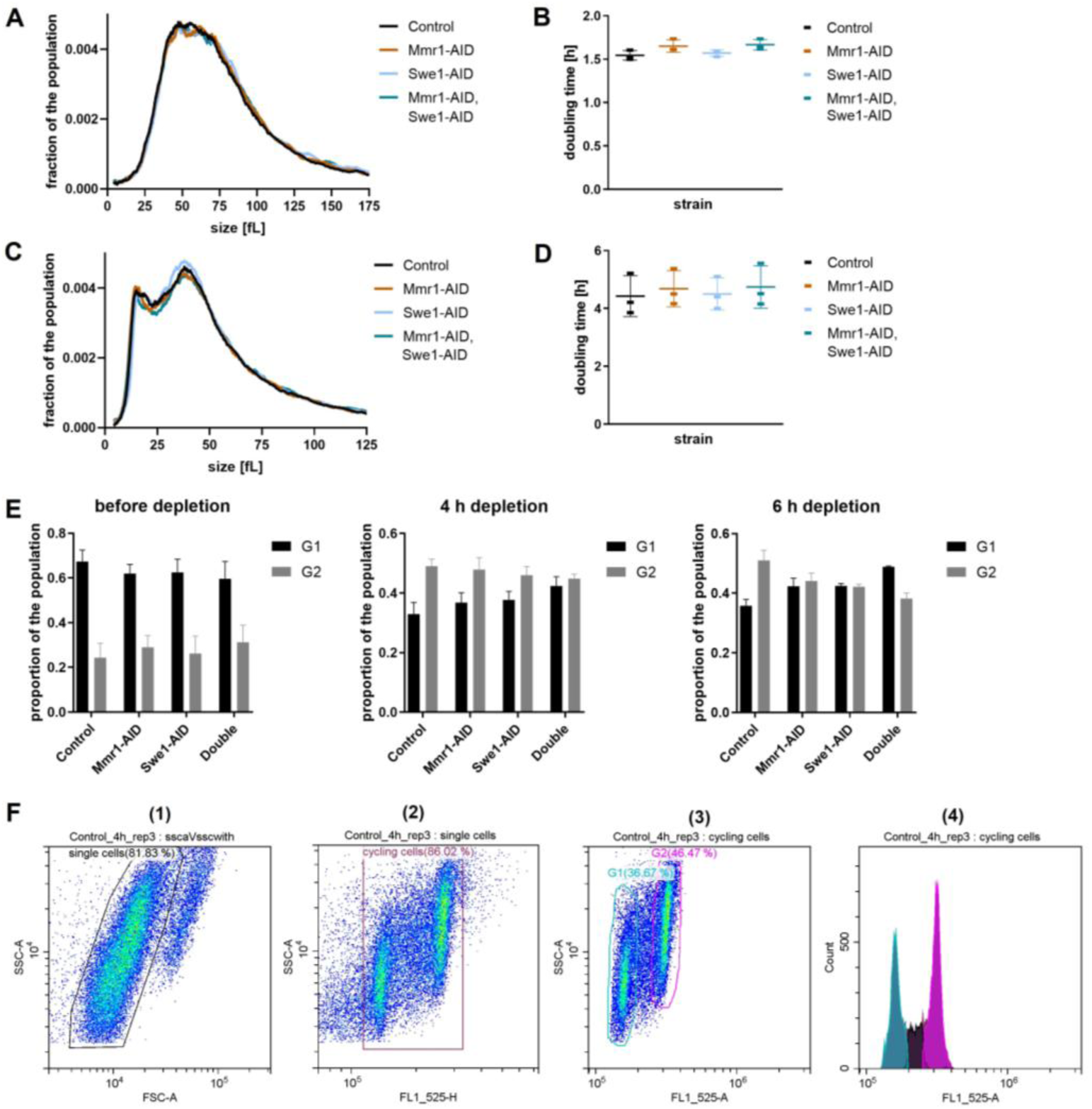
A) Cell size distribution of cells grown in SC-glucose before depletion measured using an impedance-based particle counter. The mean from three biological replicates is shown. B) Doubling times of cells grown in SC-glucose. Cells were grown to log phase and depleted using 2 µM 5-Ph-IAA. Growth was measured for 6 h after depletion and doubling times were calculated from 1-6 h after depletion. Mean and SD from three biological replicates are shown. C) Cell size distribution of cells grown in SC-ethanol before depletion measured using an impedance-based cell counter. The mean from four biological replicates is shown. B) Doubling times of cells grown in SC-ethanol. Cells were grown to log phase and depleted using 2 µM 5-Ph-IAA. Growth was measured for 6 h after depletion and doubling times were calculated from 1-10 h after depletion. Mean and SD from three biological replicates are shown. E) Proportion of cells in G1 and G2 measured by flow cytometry of SYBR green-stained cells. Mean and SD from three biological replicates are shown. F) Flow cytometry gating example. From left to right: (1) single-cells were selected (exclusion of clumps and duplets), (2) cycling cells were selected, (3) G1 and G2 cells were determined, (4) example histogram of G1, S, and G2 counts.

**Figure S5.**
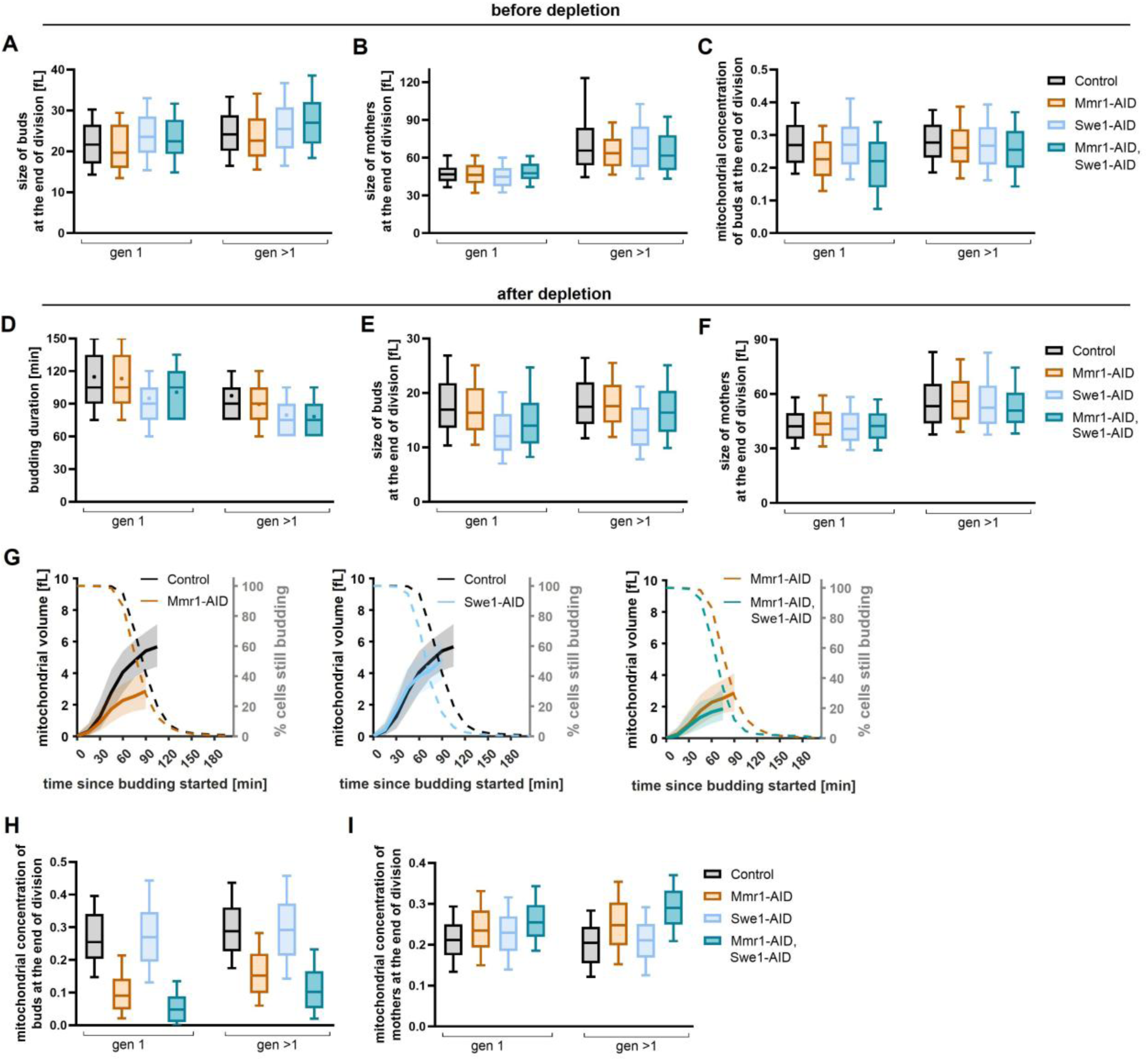
A) – I) Cells were grown in SC-ethanol and proteins were depleted through addition of 2 µM 5-Ph-IAA. Analysis from four biological replicates is shown. A-C) Analysis before depletion. Gen 1: n (Control) = 154, n (Mmr1-AID) = 118, n (Swe1-AID) = 137, n (Mmr1-AID, Swe1-AID) = 112; gen >1: n (Control) = 402, n (Mmr1-AID) = 374, n (Swe1-AID) = 325, n (Mmr1-AID, Swe1-AID) = 296. Cell sizes of buds (A) and mothers (B) at the end of division of first– and higher-generation cell cycles before depletion. C) Mitochondrial concentration of buds at the end of division for first– and higher-generation cell cycles. D) Budding duration of first– and higher-generation cell cycles after depletion. E) Cell sizes of buds of first or higher-generations mothers at the end of division after depletion. F) Sizes of firstor – higher-generation mothers at the end of division after depletion of Mmr1, Swe1, or both. G) Median mitochondrial volume (left axis, solid lines) and percentage of cells that are still budding (right axis, dashed lines) against the time since budding started. Median with 25^th^ and 75^th^ percentiles are shown for the mitochondrial volume until 85 % of the cells have finished budding. Analysis of higher-generation cell cycles is shown: n (Control) = 841, n (Mmr1-AID) = 837, n (Swe1-AID) = 1097, n (Mmr1-AID, Swe1-AID) = 893. H and I) Mitochondrial concentrations at the end of division of H) buds and I) mothers. Gen 1: n (Control) = 393, n (Mmr1-AID) = 250, n (Swe1-AID) = 417, n (Mmr1-AID, Swe1-AID) = 191; gen >1: n (Control) = 841, n (Mmr1-AID) = 837, n (Swe1-AID) = 1097, n (Mmr1-AID, Swe1-AID) = 893. For all box plots whiskers indicate 10th and 90th percentiles, line indicates median, boxes show 25th and 75th percentiles.

**Figure S6.**
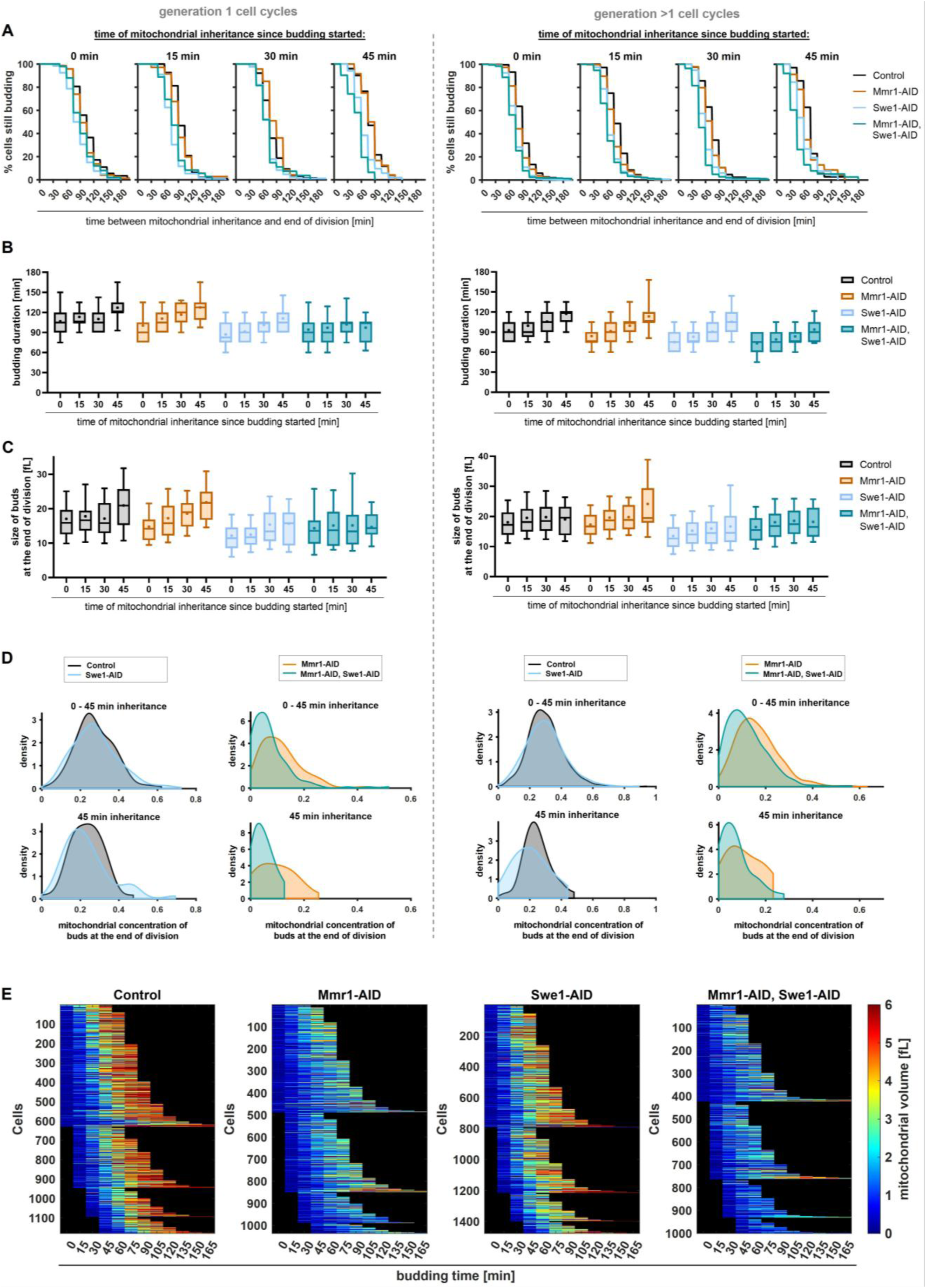
A) Time between mitochondrial inheritance and end of division. Left: first-generation, right: higher-generation cell cycles. B) Budding duration grouped by the time mitochondria are inherited of first (left) and higher-generation cell cycles (right). C) Cell size of the buds at the end of division grouped by the time mitochondria are inherited of first (left) and higher-generation cell cycles (right). B and C) Whiskers of box plots indicate 10th and 90th percentiles, line indicates median, + indicates mean, boxes show 25th and 75th percentiles. D) Half-violin plots show kernel density estimates calculated using MATLAB’s ksdensity function with zero values retained and densities area normalized. E) Heatmaps of the mitochondrial volume of buds through their budding phase. n (Control) = 1234, n (Mmr1-AID) = 1087, n (Swe1-AID) = 1514, n (Mmr1-AID, Swe1-AID) = 1084. A)-E) Cells were grown in SC-ethanol and proteins were depleted through addition of 2 µM 5-Ph-IAA. Analysis from four biological replicates is shown. A-C) Control (gen 1): n (0 min) = 145, n (15 min) = 96, n (30 min) = 64, n (45 min) = 51; Mmr1-AID (gen 1): n (0 min) = 73, n (15 min) = 71, n (30 min) = 47, n (45 min) = 24; Swe1-AID (gen 1): n (0 min) = 160, n (15 min) = 112, n (30 min) = 84, n (45 min) = 52; Mmr1-AID, Swe1-AID (gen 1): n (0 min) = 50, n (15 min) = 36, n (30 min) = 27, n (45 min) = 31. Control (gen >1): n (0 min) = 390, n (15 min) = 183, n (30 min) = 72, n (45 min) = 27; Mmr1-AID (gen >1): n (0 min) = 491, n (15 min) = 221, n (30 min) = 84, n (45 min) = 35; Swe1-AID (gen >1): n (0 min) = 647, n (15 min) = 316, n (30 min) = 99, n (45 min) = 26; Mmr1-AID, Swe1-AID (gen >1): n (0 min) = 375, n (15 min) = 308, n (30 min) = 141, n (45 min) = 38.

**Table S1.** Cell cycle genes found in the screen. Grey indicates non-essential genes, * indicates genes that were manually added since they were not assigned by KEGG to any category. ^#^ scored as essential in systematic studies.

| Category | Gene | Function | double-depleted/<br>undepleted | double-depleted/<br>single-depleted | cutoff |
| --- | --- | --- | --- | --- | --- |
| **DNA damage/**<br>**DNA replication**<br>**initiation/**<br>**replication**<br>**checkpoints** | *LCD1* | Checkpoint protein, recruits Mec1 to sites of DNA damage | 0.39 | 0.57 | 0.7 |
|  | *CDC7* | Catalytic subunit of DBF4-dependent kinase, essential for origin firing | 0.31 | 0.80 | 0.8 |
|  | *DBF4* | Regulatory subunit of CDC7 kinase, regulates replication initiation | 0.35 | 0.64 | 0.7 |
|  | *ORC5* | Subunit of Origin Recognition Complex, binds to origins of replication | 0.31 | 0.59 | 0.7 |
|  | *CDC45* | Required for initiation of DNA replication | 0.16 | 0.49 | 0.7 |
| **chromosome**<br>**condensation** | *BRN1* | Condensin complex subunit, required for chromosome condensation | 0.10 | 0.23 | 0.7 |
|  | *YCG1* | Condensin complex subunit, chromosome condensation / segregation | 0.34 | 0.66 | 0.7 |
|  | *SMC4* | Condensin complex subunit, binds chromatin and has ATPase activity | 0.37 | 0.53 | 0.7 |
| **spindle**<br>**(pole body)**<br>**components/**<br>**spindle**<br>**assembly**<br>**checkpoint** | *CIK1\** | Kinesin-associated protein, required for spindle assembly/stability | 0.74 | 0.70 | 0.8 |
|  | *DAM1* | DASH/Dam1 kinetochore complex, microtubule attachment | 0.18 | 0.71 | 0.7 |
|  | *SPC110\** | Core structural component of SPB, bridges central and inner plaques | 0.19 | 0.56 | 0.7 |
|  | *CNM67\** | Outer plaque component of SPB, required for nuclear migration | 0.49 | 0.63 | 0.7 |
|  | *SPC29\** | Inner plaque component of SPB, required for SPB duplication | 0.17 | 0.64 | 0.7 |
|  | *SPC72\** | Outer plaque component, anchors cytoplasmic microtubules to SPB, mitotic spindle orientation checkpoint | 0.63 | 0.65 | 0.7 |
|  | *SPC42\** | Central plaque component, essential for SPB duplication | 0.62 | 0.54 | 0.7 |
|  | *NBP1\** | SPB component, required for SPB duplication | 0.41 | 0.65 | 0.7 |
|  | *BUB1* | Spindle checkpoint kinase, localizes to kinetochores, delays anaphase in response to spindle and kinetochore defects | 0.63 | 0.63 | 0.7 |
|  | *BUB3* | spindle checkpoint protein in complex with Bub1 | 0.68 | 0.63 | 0.7 |
| **mitotic entry,**<br>**progression,**<br>**and exit**<br>**regulation** | *CDC15* | Protein kinase of MEN, required for mitotic exit and cytokinesis | 0.21 | 0.78 | 0.7 |
|  | *CDC5* | Polo-like kinase, regulates multiple steps of mitosis and cytokinesis | 0.44 | 0.71 | 0.8 |
|  | *ESP1* | Separase, cleaves cohesin and initiates sister chromatid separation | 0.13 | 0.77 | 0.7 |
|  | *PDS1#* | Securin, inhibits separase until anaphase onset | 0.66 | 0.70 | 0.7 |
|  | *SWE1* | Inhibits Cdc28 to prevent premature mitotic entry, G2/M checkpoint | 0.79 | 0.74 | 0.8 |
|  | *APC1* | APC subunit, essential for cyclin degradation and mitotic exit | 0.25 | 0.54 | 0.7 |
|  | *APC4* | APC subunit, essential for cyclin degradation and mitotic exit | 0.44 | 0.62 | 0.7 |
|  | *APC5* | APC subunit, essential for cyclin degradation and mitotic exit | 0.25 | 0.44 | 0.7 |
|  | *CDC16* | Tetratricopeptide repeat (TPR) subunit of the APC | 0.76 | 0.69 | 0.8 |
|  | *CDC27* | TPR subunit of the APC | 0.16 | 0.37 | 0.7 |
|  | *DOC1* | APC subunit, involved in substrate recognition | 0.65 | 0.71 | 0.7 |
| **others** | *CIP1\** | Cyclin-dependent kinase inhibitor, regulates G1/S transition | 0.46 | 0.44 | 0.7 |
|  | *FAR7\** | Component of Far complex at ER | 0.77 | 0.80 | 0.8 |
|  | *PHO5* | Acid phosphatase | 0.80 | 0.71 | 0.8 |
|  | *SOG2\** | RAM network component, regulates cell morphogenesis and separation | 0.53 | 0.62 | 0.7 |

**Table S2.** Spliceosome genes from the screen. Grey indicates non-essential genes, * indicates genes that were manually added since they were not assigned by KEGG to any category.

| Gene | Function | double-depleted/<br>undepleted | double-depleted/<br>single-depleted | cutoff |
| --- | --- | --- | --- | --- |
| *BUD31* | Spliceosome assembly and catalytic activity | 0.67 | 0.72 | 0.8 |
| *LSM3* | U6 snRNA-binding, tri-snRNP formation | 0.56 | 0.64 | 0.7 |
| *LSM5* | U6 snRNA-associated Sm-like protein | 0.48 | 0.50 | 0.7 |
| *PRP16* | ATPase, splicing fidelity and proofreading | 0.73 | 0.72 | 0.8 |
| *PRP18* | Catalytic step II in splicing | 0.49 | 0.70 | 0.7 |
| *PRP2* | Spliceosome ATPase, splicing activation | 0.27 | 0.51 | 0.7 |
| *PRP22* | RNA helicase, mRNA release from spliceosome | 0.25 | 0.73 | 0.8 |
| *PRP3* | U4/U6 di-snRNP, tri-snRNP assembly | 0.21 | 0.71 | 0.8 |
| *PRP45* | Essential spliceosome-associated protein | 0.13 | 0.23 | 0.7 |
| *PRP46* | Essential spliceosome-associated protein | 0.41 | 0.72 | 0.8 |
| *PRP5* | DEAD-box helicase, prespliceosome formation | 0.10 | 0.44 | 0.7 |
| *PRP6* | U4/U6-U5 tri-snRNP complex component | 0.12 | 0.69 | 0.7 |
| *SLU7* | 3' splice site selection, gene regulation | 0.21 | 0.77 | 0.8 |
| *SMB1* | Core Sm protein, snRNP component | 0.29 | 0.25 | 0.7 |
| *SNP1* | U1 snRNP protein, 5' splice site recognition | 0.21 | 0.70 | 0.7 |
| *SNU66* | Tri-snRNP component, Hub1 interaction | 0.73 | 0.78 | 0.8 |
| *SYF1* | Splicing, transcription elongation, TREX complex | 0.31 | 0.48 | 0.7 |
| *THO2* | mRNA export, transcription elongation | 0.31 | 0.39 | 0.7 |
| *YHC1* | U1 snRNP, 5' splice site binding | 0.20 | 0.77 | 0.8 |
| *SMX2* | Core Sm protein Sm G, part of the spliceosomal snRNPs | 0.13 | 0.33 | 0.7 |
| *SNU56\** | U1 snRNP, meiosis-specific splicing | 0.20 | 0.51 | 0.7 |
| *LUC7\** | U1 snRNP, 5' splice site recognition | 0.16 | 0.70 | 0.7 |
| *PRP24\** | U6 snRNA chaperone, U4/U6 annealing | 0.00 | 0.00 | 0.7 |
| *CWC25\** | Step 1 splicing factor, spliceosome binding | 0.35 | 0.65 | 0.7 |

**Table S3.** Information about respiratory deficiency/petite/mtDNA of the cell cycle genes which were found as negative genetic interactors in our screen.

| Gene | Petite/mtDNA loss/respiratory deficient |
| --- | --- |
| *DOC1* | pet gene, class II pet gene (Stenger, Le et al. 2020)<br>pet gene class I (required for mtDNA maintenance (Merz and Westermann 2009)<br>no mtDNA (Göke, Schrott et al. 2020)<br>0.1x mtDNA levels (Puddu, Herzog et al. 2019)<br>respiratory deficient (Luban, Beutel et al. 2005) |
| *CNM67* | pet gene (Stenger, Le et al. 2020) |
| *SPC72* | respiratory deficient (Luban, Beutel et al. 2005)<br>pet gene (Stenger, Le et al. 2020)<br>increased mtDNA (2.42x, qPCR, relative cell size 2.64 (Göke, Schrott et al. 2020))<br>increased mtDNA (33x (Puddu, Herzog et al. 2019))<br>pet gene (Dimmer, Fritz et al. 2002)<br>small colonies (Soues and Adams 1998)<br>pet gene (Merz and Westermann 2009) |
| *BUB1* | slow growth on glycerol medium (Dimmer, Fritz et al. 2002) |
| *BUB3* | pet gene (Stenger, Le et al. 2020)<br>increased mtDNA (2.67x, relative cell size 1.48 (Göke, Schrott et al. 2020))<br>slow growth on glycerol medium (Dimmer, Fritz et al. 2002) |

**Table S4.**
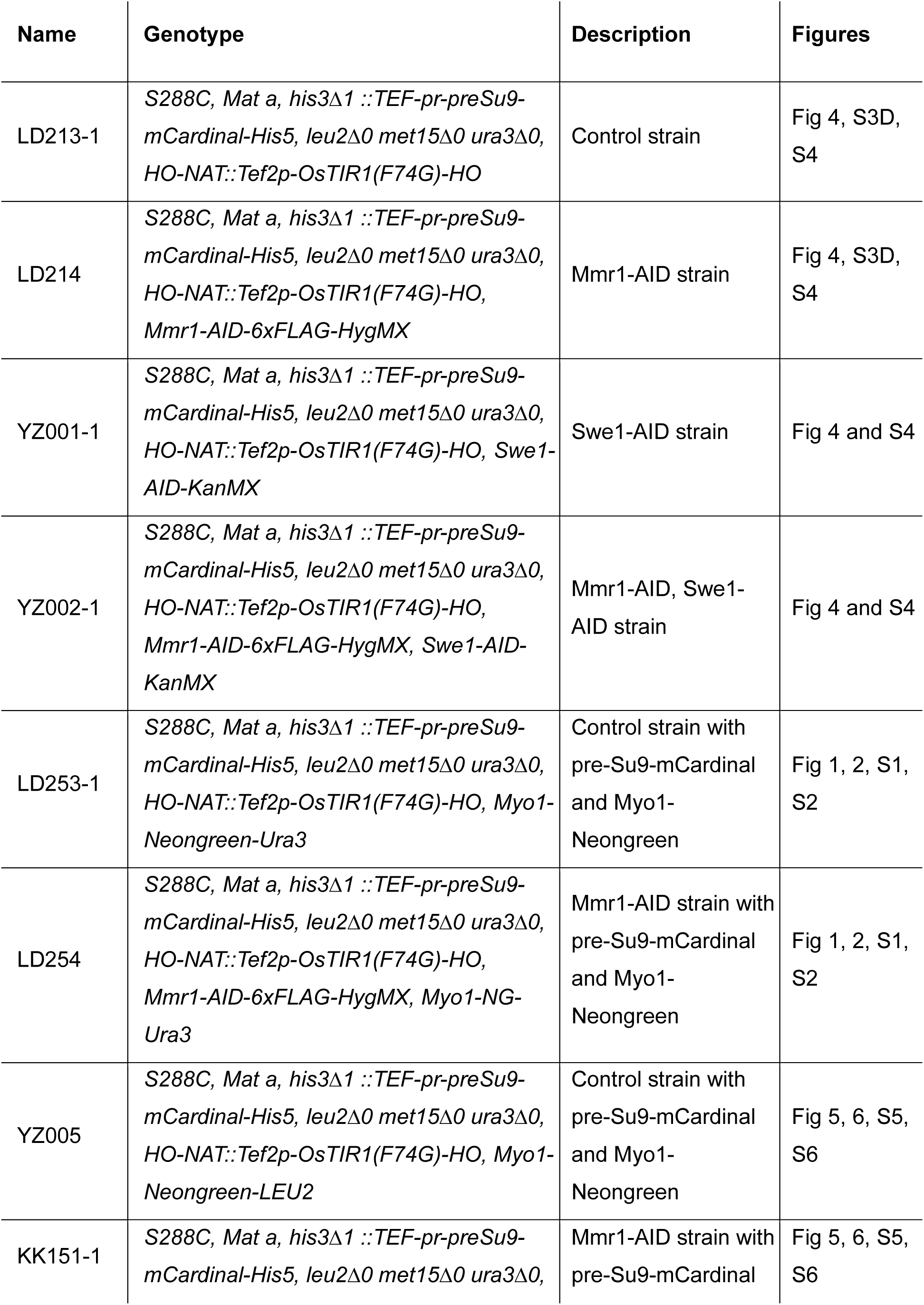
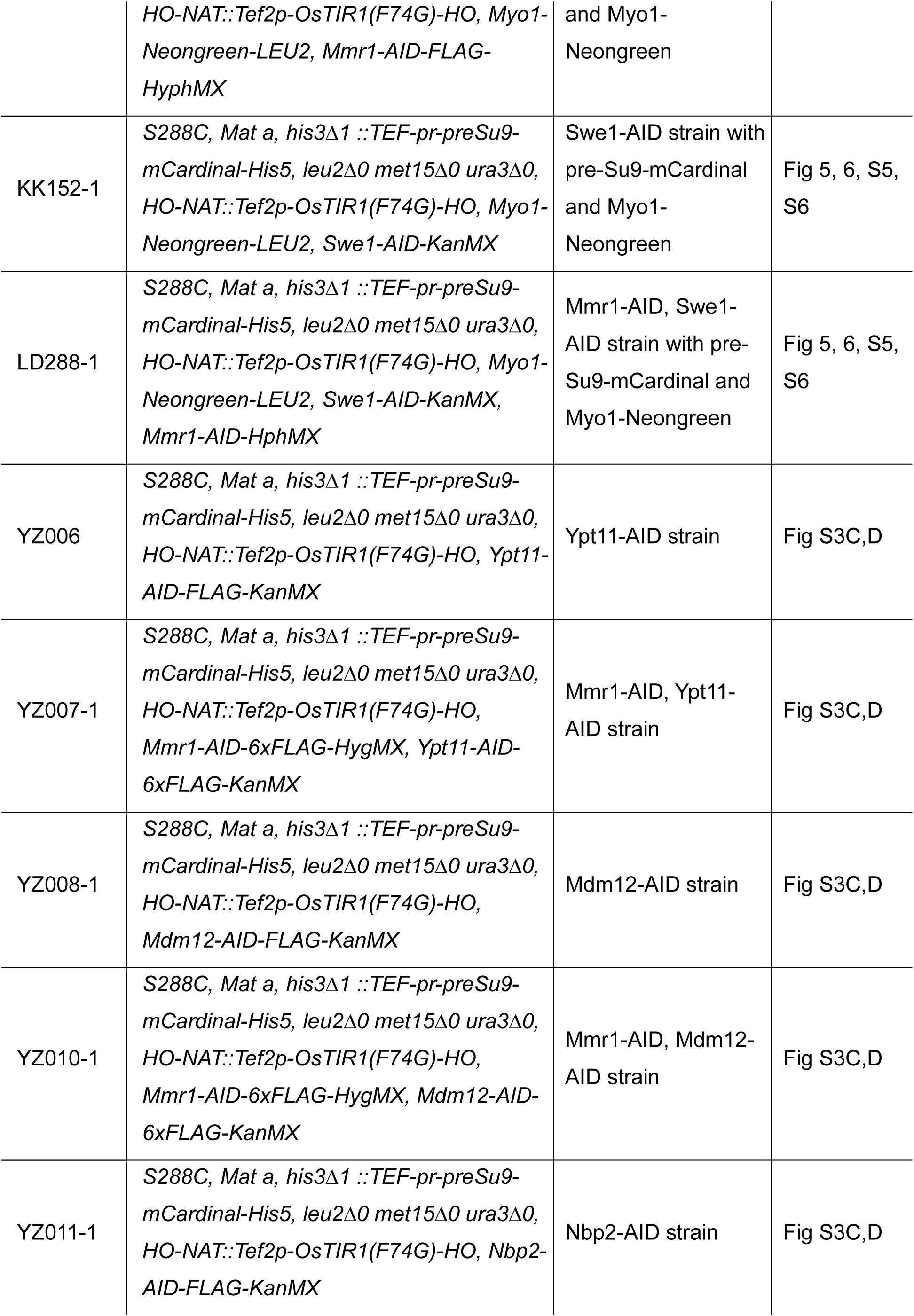
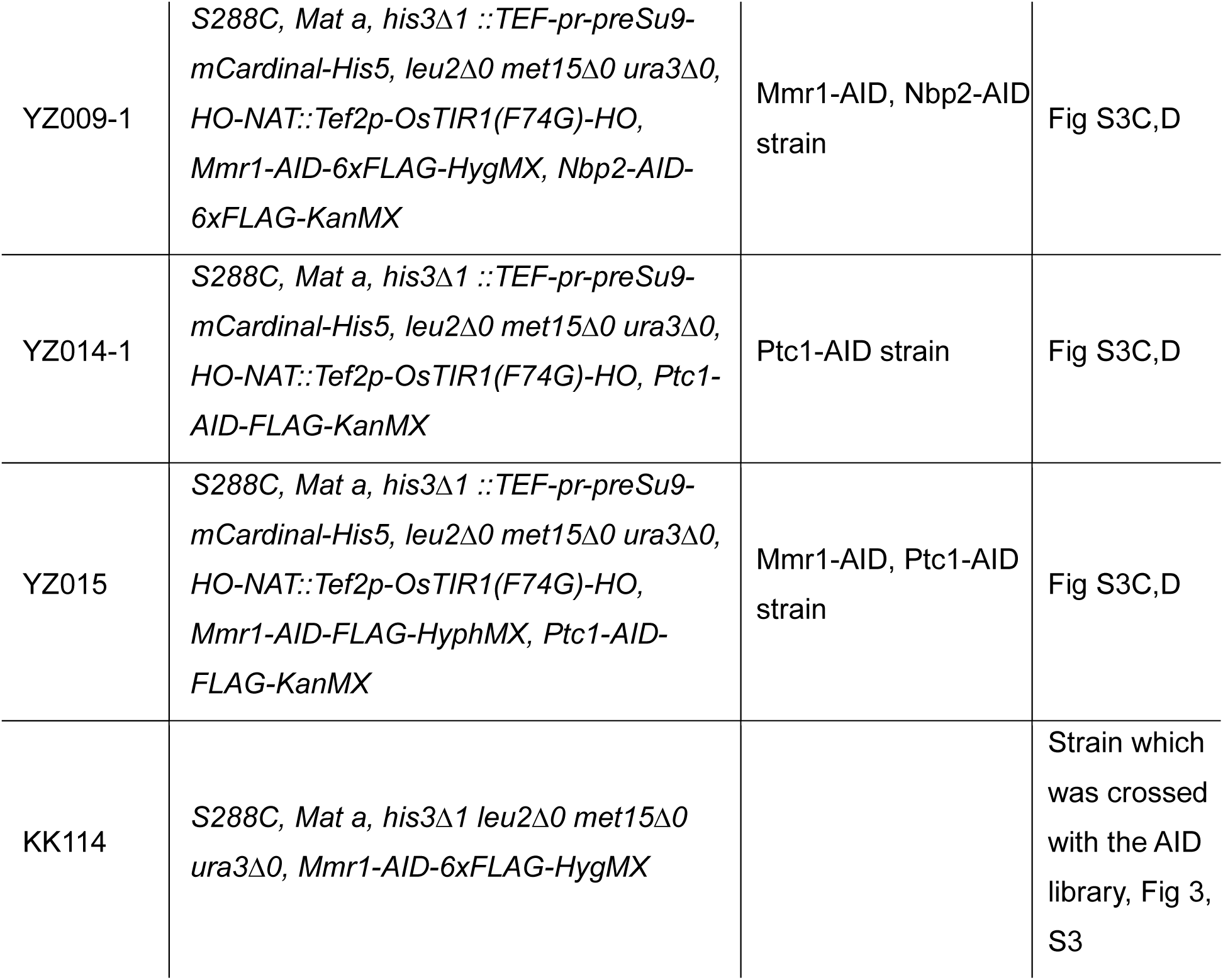
Strains used in this study. All strains are By4741 and BY4742 derivatives. All strains were constructed in this study.

**Table S5.** Plasmids used in this study.

| Name | Description | Origin |
| --- | --- | --- |
| pLD036 | TEFpr-preSu9-mCardinal-His5 | This study |
|  | *AID-6xFLAG-HygMX* | gift from Helle Ulrich (Addgene plasmid #99519;<br>http://n2t.net/addgene:99519;RRID:Addgene_99519)<br>(Morawska and Ulrich 2013) |
| pKK029 | *AID-6xFLAG-KanMX* | This study |

**Supplementary Table S6.** Components of 1x Synthetic complete (SC).

| component | g/L | [mM] final |
| --- | --- | --- |
| Adenine | 0.031 | 0.228 |
| L-Arg (HCl) | 0.021 | 0.098 |
| L-Aspartic acid | 0.103 | 0.773 |
| L-Glutamic acid | 0.081 | 0.548 |
| L-His | 0.021 | 0.133 |
| L-Leu | 0.123 | 0.941 |
| L-Lys (HCl) | 0.031 | 0.169 |
| L-Met | 0.021 | 0.138 |
| L-Phe | 0.051 | 0.311 |
| L-Ser | 0.386 | 3.670 |
| L-Thr | 0.206 | 1.727 |
| L-Tyr | 0.031 | 0.170 |
| L-Trp | 0.041 | 0.201 |
| L-Val | 0.154 | 1.317 |
| Uracil | 0.021 | 0.184 |

**TableS7. Optical.** filters for epifluorescence microscopy. Filters used for imaging on a Nikon Ti2-E epifluorescence microscope. All described filters are manufactured by Chroma and purchased from AHF.

| Fluorophore | LED Wave-length | Filter Set | Excitation Filter | Dichroic | Emission Filter | Figures |
| --- | --- | --- | --- | --- | --- | --- |
| mNeongreen | 513 nm | YFP ET Filter Set | ET500/20x | T515lp, Di 25 mm x 36 mm | ET535/30m | Figure 1, 2, S1, S2 |
| mCardinal | 575 nm | 585/29 BrightLine HC 650/60 BrightLine HC Beamsplitter T610 LPXR | ET585/29x | T610 LPXR, Di 25 mm x 36 mm | ET650/60 | Figure 1, 2, S1, S2 |
| mNeongreen | 508 nm | RNA mango: TO1-set HC BS 518 HC 535/22 | BrightLine Fluorescence Filter 504/12 | HC BS 518 | HC 535/22 | Figure 5, 6, S5, S6 |
| mCardinal | 555 nm | Set far-red HC BS 660 HC 692/40 | BrightLine Fluorescence Filter 556/20 | HC BS 660 | HC 692/40 | Figure 5, 6, S5, S6 |

**TableS8. Exposure.** times and intensities for epifluorescence microscopy.

| Fluorophore | Imaged protein | Intensity | Exposure time | Figures |
| --- | --- | --- | --- | --- |
| mCardinal | preSu9 | 5 % | 100 ms | Figure 1, 2, S1, S2 |
| mNeongreen | Myo1 | 10 % | 200 ms | Figure 1, 2, S1, S2 |
| mCardinal | preSu9 | 40 % | 300 ms | Figure 5, 6, S5, S6 |
| mNeongreen | Myo1 | 20 % | 300 ms | Figure 5, 6, S5, S6 |

